# Active Vertex Model for Cell-Resolution Description of Epithelial Tissue Mechanics

**DOI:** 10.1101/095133

**Authors:** Daniel L. Barton, Silke Henkes, Cornelis J. Weijer, Rastko Sknepnek

## Abstract

We introduce an Active Vertex Model (AVM) for cell-resolution studies of the mechanics of confluent epithelial tissues consisting of tens of thousands of cells, with a level of detail inaccessible to similar methods. The AVM combines the Vertex Model for confluent epithelial tissues with active matter dynamics. This introduces a natural description of the cell motion and accounts for motion patterns observed on multiple scales. Furthermore, cell contacts are generated dynamically from positions of cell centres. This not only enables efficient numerical implementation, but provides a natural description of the T1 transition events responsible for local tissue rearrangements. The AVM also includes cell alignment, cell-specific mechanical properties, cell growth, division and apoptosis. In addition, the AVM introduces a flexible, dynamically changing boundary of the epithelial sheet allowing for studies of phenomena such as the fingering instability or wound healing. We illustrate these capabilities with a number of case studies.

## I. INTRODUCTION

Collective cell migration^1,2^ in epithelial tissues is one of the key mechanisms behind many biological processes, such as the development of an embryo,^3^ wound healing,^4,5^ and tumour metastasis and invasion.^6^ Due to their layered, tightly connected structure,^7^ epithelial tissues also serve as an excellent model system to study cell migration processes. Over several decades^8^ extensive research efforts have been devoted to understanding molecular processes that lead to cell migration^9^ and, at larger scales, on how cell migration drives complex processes at the level of the entire tissue, such as morphogenesis. With recent advances in various microscopy techniques combined with the development of sophisticated automatic cell tracking methods, it is now possible to study collective migration patterns of a large number of cells over extended periods of time with the cell-level resolution, both *in vitro* and *in vivo.* Traction force microscopy^10^ experiments revealed that collective cell motion is far more complex than expected.^11–13^It is often useful to draw parallels between the collective behaviour of tissues and systems studied in the physics of colloids, granular materials and foams as these can provide powerful tools for understanding complex cell interactions in biological systems. For example, in a homogenous cell monolayer, one observes large spatial and temporal fluctuations of inter-cellular forces that cannot be pinpointed to a specific location, but cover regions that extend over several cells.^14^ These are reminiscent of the fluctuations observed in supercooled colloidal and molecular liquids approaching the glass transition^11^ and include characteristic features of the dynamical and mechanical response, such as dynamical heterogeneities and heterogeneous stress patterns, that were first observed in glasses, colloids and granular materials and that have extensively been studied in soft condensed matter physics.^15^ It has also been argued that the migration patterns are sensitive to the expression of different adhesion proteins^16^ as well as to the properties of the extracellular environment,^17^ such as the stiffness of the substrate.^18,19^ These observations lead to the development of the notion of *plithotaxis*,^12^ a universal mechanical principle akin to the more familiar chemotaxis, which states that each cell tends to move in a way that maintains minimal local intercellular shear stress. While plausible, it is yet to be determined whether plithotaxis is indeed a generic feature in all epithelial tissues.

Equally fascinating are the experiments on model systems that study cell migration in settings designed to mimic wound healing.^5,20–23^ For example, the existence of mechanical waves that span the entire tissue and generate long-range cell-guidance have been established in Madin-Darby Canine Kidney (MDCK) epithelial cell monolayers.^23^ Subtle correlations between purse-string contractility and large-scale remodelling of the tissue while closing circular gaps have also been identified.^22^ Finally, a mechanism dubbed *kenotaxis* has been proposed,^20^ which suggests that there is a robust tendency of a collection of migrating cells to generate local tractions that systematically and ooperatively pull towards the empty regions of the substrate.

On the developmental side, in pioneering work, Keller *et al.*^24^constructed a light-sheet microscope that enabled them to track *in vivo* positions of each individual cell in a zebrafish embryo over a period of 24h. A quantitive analysis^25^ of the zebrafish embryo was also able to relate mechanical energy and geometry to the shapes of the aggregate surface cells. Another extensively studied system that allows detailed tracking of individual cells is the *Drosophila* embryo.^26–30^ In recent studies that combined experiments with advanced data analysis, it was possible to quantitatively account for shape change of the wing blade by decomposing it into cell divisions, cell rearrangements and cell shape changes.^31,32^ Finally, it has recently become possible to track more than 100,000 individual cells in a chick embryo over a period exceeding 24 hours.^33^ This was achieved by developing an advanced light-sheet microscope and state-of-the-art data analysis techniques designed to automatically track individual cells in a transgenic chick embryo line with the cell membranes of all cells in the embryonic and extra embryonic tissues labelled with a green fluorescent protein tag. All these experiments and advanced data analysis techniques provide unprecedented insights into the early stages of the embryonic development, making it possible to connect processes at the level of individual cells with embryo-scale collective cell motion patterns.

While there have been great advances in our understanding of how cells regulate force generation and transmission between each other and with the extracellular matrix in order to control their shape and cell-cell contacts,^9^ it is still not clear how these processes are coordinated at the tissue-level to drive tissue morphogenesis or allow the tissue to maintain its function once it reaches homeostasis. Computer models of various levels of complexity have played an essential role in helping to answer many of those questions. ^34^ One of the early yet successful approaches has been based on the cellular Potts model (CPM).^35,36^ In the CPM, cells are represented as groups of “pixels” of a given type. Pixels are updated one at a time following a set of probabilistic rules. Pixel updates in the CPM are computationally inexpensive,^37^ which allows for simulations of large systems. In addition, the CPM extends naturally to three dimensions.^38^ While very successful in describing cell sorting as well as certain aspects of tumour growth,^39^ CPM has several limitations, the main one being that the dynamics of pixel updates is somewhat artificial and very hard to relate to the dynamics of actual cells. This problem has been to some extent alleviated by the introduction of the subcellular element model (ScEM).^40,41^ ScEM is an off-lattice model with each cell being represented as a collection of 100-200 elements - spherical particles interacting with their immediate neighbours via short-range soft-core potentials. Therefore, ScEM is able to model cells of arbitrary shapes that grow, divide and move in 3D. The main disadvantage of ScEM is that it is computationally expensive (off-lattice methods in general require more computations per time step compared to their lattice counterparts), and without a highly optimised parallel implementation, applications of the ScEM are limited to a few hundred cells at most, which is not sufficient to study effects that span long length- and time-scales.

Particle-based models have also been very successful at capturing many aspects of cell migration in tissues.^42–44^ However, when it comes to modelling confluent epithelial layers with the resolution of individual cells, one of the most widely and successfully used models is the Vertex Model (VM). The VM originated in studies of the physics of foams in the 1970s and was first applied to model monolayer cell sheets in the early 1980s.^45^ Over the past 35 years it has been implemented and extended numerous times and used to study a wide variety of different systems.^46–51^ The VM is in the core of the cell-based^52^ CHASTE,^53^ a versatile and widely used software package. The VM assumes that all cells in the epithelium are roughly the same height and thus that the entire system can be well approximated as a two-dimensional sheet. The conformation of the tissue in the VM is computed as a configuration that simultaneously optimises area and perimeter of all cells. While the model is quasi-static in nature, it captures remarkably well many properties of actual epithelial tissues. There have been numerous attempts to introduce dynamics into the vertex model.^45,49,51,55,56^ However, there are limitations associated with each approach. To the best of our knowledge, most dynamical versions of the vertex model seem to neglect fluctuations with a notable recent exception.^51^ In contrast, recent traction microscopy experiments^14^ suggest that these fluctuations might be a crucial ingredient in understanding collective cell migration. Finally, we point out a technical point that makes the implementation of VM somewhat challenging. Namely, in order to capture topology changing moves, such as cell neighbour exchanges, i.e., T1 transitions, one has to perform rather complex mesh restructuring operations^49,54^ that require complex data structures and algorithms and that inevitably add to the computational complexity of the model.

Building upon the recently introduced Self-Propelled Voronoi (SPV) model,^57^ in this paper we apply the ideas introduced in the physics of active matter systems^58^ to the VM. This allows us to construct a hybrid, Active Vertex Model (AVM) that is able to accurately describe the collective migration dynamics of a large number of cells. The AVM is implemented within *SAMoS,* an off-lattice particle-based simulation software developed specifically to study active matter systems. One of the key advantages of the hybrid approach presented in this study is that it not only enables studies of very large systems, but also introduces a very natural way to handle the T1 transitions, thus removing the need for complex mesh manipulations that are of uncertain physical and biological meaning.

The paper is organised as follows. In Section II we briefly review the general features of the VM and derive expressions for forces on individual cells central to the active vertex model. In Section III we apply the model to a number of test cases. Finally, in Section IV we provide an outlook for future applications of this model. Details of the force calculation and implementation are presented in the Appendices.

## II. MODEL

### A. Overview of the Vertex Model

Owing to its origins in the physics of foams,^59^ in the VM cells are modelled as two-dimensional convex polygons that cover the plane with no holes and overlaps, i.e., the epithelial tissue is represented as a convex polygonal partitioning of the plane (Fig. 1). The main simplification compared to models of foams is that the VM assumes that contacts between neighbouring cells are straight lines. In addition, neighbouring cells are also assumed to share a single edge, which is a simplification compared to real tissues, where junctions between two neighbouring cells consist of two separate cell membranes that can be independently regulated. Typically, three junction lines meet at a vertex, although vertices with a higher number of contacts are also possible.^54^ The model tissue is therefore a mesh consisting of polygons (i.e., cells), edges (i.e., cell junctions), and vertices (i.e., meeting points of three or more cells).

**Figure 1.**
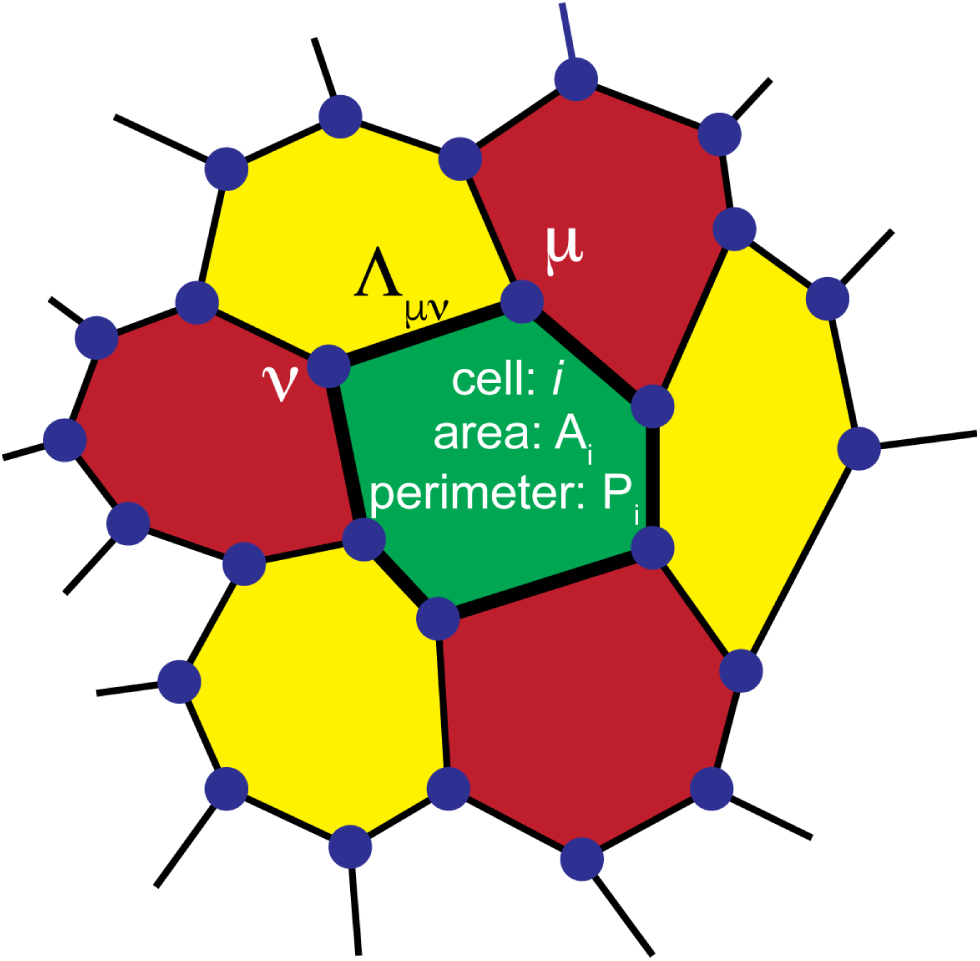
In the Vertex Model (VM), a confluent epithelial sheet is represented as a polygonal tiling of the plane with no holes or overlaps. Each cell is represented by an *n* — sided polygon. Neighbouring cells share an edge, which models the cell junction as a straight line. Three edges meet at a vertex (dark blue dots). The behaviour of cell *i* is described by three parameters: 1) reference area 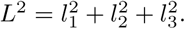, 2) area modulus *K_i_*, and 3) perimeter modulus Γ*_i_*. In addition, a junction connecting vertices *μ* and *ν* has tension *Λ μ* ν.

An energy is associated to each configuration of the mesh. It can be written as

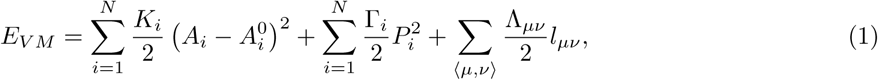

where *N* is the total number of cells, *A_i_* is the area of the cell *i*, while 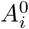 is its reference area. *K_i_* is the area modulus, i.e. a constant with units of energy per area squared measuring how hard it is to change the cell’s area. *P_i_* is the cell perimeter and Γ*_i_* (with units of energy per length squared) is the perimeter modulus that determines how hard it is to change perimeter *P_i_*. *l_μν_* is the length of the junction between vertices *μ* and *ν* and Λ*_μν_* is the tension of that junction (with units of energy per length). 〈*μ*, *ν*〉 in the last term denotes the sum is over all pairs of vertices that share a junction. Note that the model allows for different cells to have different area and perimeter moduli as well as reference areas, allowing for modelling of tissues containing different cell types.

Some authors write the perimeter term as 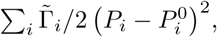, where 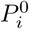 is a reference perimeter, and omit the last term in Eq. (1) or completely omit the *P*^2^ term.^45,54^ Under the assumption that the values of Λ*_μν_* for all junctions of the cell i are the same, i.e., Λ*μ*,*ν* ≡ Λ*_i_*, the last term in Eq. (1) becomes 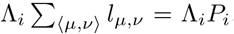. Therefore, if we identify 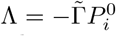, it immediately becomes clear that the descriptions in Eq. (1) and the model with the preferred perimeter are identical to each other. Note that this is true up to the constant term 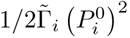, which is unimportant as it only shifts the overall energy and does not contribute to the force on the cell (see below). The description in Eq. (1) is slightly more general as it allows for the junctions to have different properties depending on the types of cells that are in contact. Retaining the 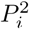 term is also advisable in order to prevent the model from becoming unstable if the area modulus is too small.

It is straightforward to express cell area and cell perimeter in terms of vertex coordinates. Therefore, vertex positions together with their connectivities uniquely determine the energy of the epithelial sheet, hence the name Vertex model. The main assumption of the VM is that the tissue will always be in a configuration which minimises the total energy in Eq. (1). Determining the minimum energy configuration is a non-trivial multidimensional optimisation problem and, with the exception of a few very simple cases, it can only be solved numerically. A basic implementation of the VM therefore needs to use advanced multidimensional numerical minimisation algorithms to determine the positions of vertices that minimise Eq. (1) for a given set of parameters *K_i_*, Γ*_i_* and Λ*_μν_*. Most implementations,^49,51,54^ also introduce topology (connectivity) changing moves to model events such as cell rearrangements.

There have been several attempts to introduce dynamics into the VM,^45,49,55^ including a recent study that introduced stochasticity into the junction tension.^51^ The idea behind such approaches it to write equations of motion for each vertex as

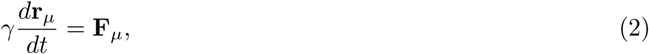

where *γ* is a friction coefficient and Γ*_μ_* is the position vector of vertex *μ*. F*_μ_* is the total force on vertex *μ* computed as the negative gradient of Eq. (1) with respect to **r***_μ_*, i.e., **F**_*μ*_ = –∇_r_*μ*__*E_V M_*. We note that the exact meaning of friction in confluent epithelial tissues is the subject of an ongoing debate that is beyond the scope of this study. Here, as in the case of most models to date, we assume that all effects of friction (i.e., between neighbouring cells as well as between cells and the substrate and the extracellular matrix) can be modelled by a single constant. While this may appear to be a major simplification, as we will show below, the model is capable of capturing many key features of real epithelial tissues. Eq. (2) is a first order equation since the mass terms have been omitted. This so-called overdamped limit is very common in biological systems, since the inertial effects are typically several orders of magnitude smaller than the effects arising from the cell-cell interactions or random fluctuations produced by the environment. Note that the force on vertex *μ* depends on the position of its neighbouring vertices, resulting in a set of coupled non-linear ordinary differential equations. In most cases those equations can only be solved numerically.

While the introduction of dynamics alleviates some of the problems related to the quasi-static nature of the VM, one still has to implement topology changing moves if the model is to be applicable to describing cell intercalation events. This can lead to unphysical back and forth flips of the same junction and has only recently been analysed in full detail.^54^

### B. Active Vertex Model

It is important to note that the VM in its original form is a *quasi-static* model. In other words, it assumes that at every instant in time, the tissue is in a state of mechanical equilibrium. This is a strong assumption, which is in line with many biological systems, especially in the case of embryos where cells actively grow, divide and rearrange. As a matter of fact, biological systems are among the most common examples of systems out of equilibrium. Therefore, while it is able to capture many of the mechanical properties of the tissue, the VM is unable to fully describe the effects that are inherently related of being out of equilibrium. Many such effects are believed to be behind the collective migration patterns observed in recent experiments. In addition, in many dynamical implementations of the VM the effects of both thermal and non-thermal random fluctuations originating in complex intercellular processes and interactions with the environment are either completely omitted or not very clear. While for a system out of equilibrium the fluctuation-dissipation theorem^61^ does not hold, and the relation between random fluctuations and friction is not simple, it is even more important to note that fluctuations can have non-trivial effects on the collective motion patterns.^51^

Here we take an alternative approach and build a description similar to the recently introduced SPV model.^57^The idea behind the SPV is that instead of treating vertices as the degrees of freedom, one tracks positions of cell centres. Forces on cell centres are, however, computed from the energy of the VM, Eq. (1). The core assumption of the model is that the tissue confirmations correspond to the Voronoi tessellations of the plane with cell centres acting as Voronoi seeds. We recall that a Voronoi tessellation is a polygonal tiling of the plane based on distances to a set of points, called *seeds.* For each seed point there is a corresponding polygon consisting of all points closer to that seed point than to any other. This imposes some restrictions onto possible tissue confirmations, i.e., not all convex polygonal tessellations of the plane are necessarily Voronoi, but it has recently been argued that Voronoi tessellations can predict the diverse cell shape distributions of many tissues.^62^ Furthermore, the exact details of the tessellation are not expected to play a significant role in the large scale behaviour of the tissue, which this model aims to describe.

In the original implementation of Bi, *et al.*^57^the Voronoi tessellation is computed at every time step. The vertices of the tessellation are then used to evaluate forces at all cell centres, that are, in turn, moved in accordance to those forces and the entire process is repeated. While conceptually clear, this procedure is numerically expensive as it requires computation of the entire Voronoi diagram at each time step. This limits the accessible system size to several hundred cells.

Here, we instead propose an alternative approach based on the Delaunay triangulation. The Delaunay triangulation for a set of points *P* in the plane is a triangular tiling, *DT* (*P*), of the plane with the property that there are no points of *P* inside the circumcircle of any of the triangles in *DT* (*P*).^63^ A property of a Delaunay triangulation that is key for this work is that it is possible to construct a so-called dual Voronoi tessellation by connecting circumcenters of its triangles. This establishes a mathematical duality between Delaunay and Voronoi descriptions. This duality is exact and quantities, such as the force, expressed on the Voronoi tiling have an exact map onto quantities expressed on its dual Delaunay triangulation. Although being non-linear (see Sec. IIB 1), this map is relatively simple, and therefore fast to compute. An important property of the Voronoi-Delaunay duality is that continuous deformations of one map into continuous deformations of the other. In other words, smooth motion of a cell’s centre will correspond to a smooth change in that cell’s shape. This is crucial to ensuring that during the dynamical evolution the cell connectivity changes continuously, a feature that is essential for accurately modelling T1 transitions. The main advantage of working with the Delaunay description is that while the Voronoi tessellation has to be recomputed each time cell centres move, it is possible to retain the Delaunay character of a triangulation via local edge flip moves (Fig. 2c), which drastically increases the efficiency of the Delaunay based approach and enables us to simulate systems containing tens of thousands of cells.

**Figure 2.**
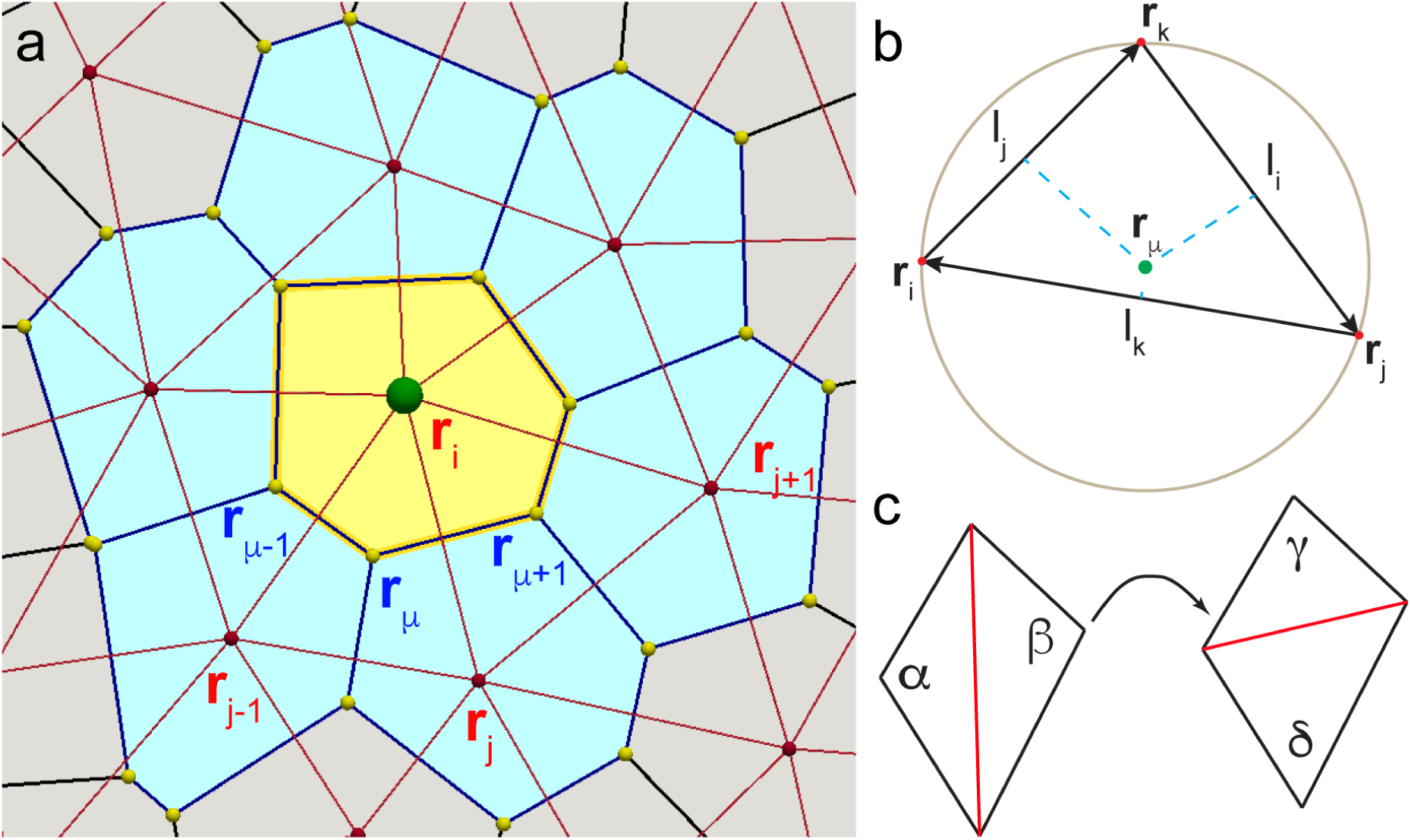
a) Two coordinate representations dual to each other. In the AVM we track particles that correspond to cell centres (red/green dots). Particles form a Delaunay triangulation (red lines) and their positions are labeled with vectors carrying Latin indices. The dual of the Delaunay triangulation is a Voronoi tessellation, with each Voronoi cell (a shaded polygon) representing an actual cell. Cell edges are marked by blue lines. Positions of the vertices (yellow dots) of the dual mesh are denoted by Greek indices. b) Position of the circumcenter **r***_μ_* of the triangle with corners **r***_i_* **r***_j_* and **r***_k_*. c) The edge flip move is at the core of the equiangulation procedure. If the sum of the angles opposite to the red edge exceeds *α* + *β* > 180° the edge is "flipped". As the result, the sum of the angles opposite to the new edge is less than 180°, i.e., *γ* + *δ* < 180°. Note that this edge flip is local and only affects one Voronoi edge (i.e., cell junction). Therefore, edge flips can only affect the local connectivity of four cells (see also, Fig. 5).

Before we introduce the Active Vertex Model (AVM), we pause to make a comment about the notation. In the following, we will always use Latin letters to denote cells, i.e. positions of their centres, and Greek letters to denote vertices of the dual Voronoi tessellation, i.e., meeting points of three or more cells. Therefore, vertices of the VM will always carry Greek indices (Fig. 2a).

### 1. Force on the cell centre

In the AVM, the area *A_i_* in Eq. (1) corresponding to the cell (particle) *i* is the area of the polygon (face) of the Voronoi polygon, Ω*_i_*, associated with *i*. It is given as

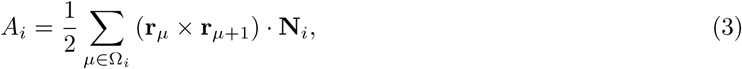

where **r***_μ_* is the position of vertex *μ* and **N**_*i*_ ≡ e*_z_* is a unit-length vector perpendicular to the surface of the polygon - which does not depend on the position of the vertices in the plane. The sum is over all vertices of the Voronoi cell and we close the loop with *μ* +1 ≡ 1 for *μ* = *N*_Ω_*_i_*, where *N*_Ω_*_i_* is the total number of vertices in the cell Ω*i*. This expression is just a discrete version of Green’s formula. Similarly, the cell perimeter is

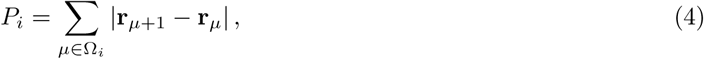

with the same rules for the next neighbour of the “last” vertex. |·| represents the Euclidean length of the vector.

In order to make a connection between cell centres and indices of the Voronoi tessellation we recall that for a given triangle in the Delaunay triangulation, the position of its dual Voronoi vertex coincides with the centre of the circumscribed circle. The position of the circumcenter is given by (see Fig. 2b)

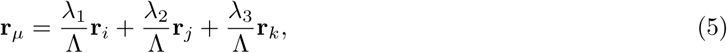

where **r***_i_*, **r***_j_* and **r***_k_* are position vectors of the corners of the triangle and λ_1_, λ_2_ and λ_3_are the barycentric coordinates (see, Eq. (26) in Appendix I), with Λ = λ_1_ + λ_2_+ λ_3_.

Armed with the mapping in Eq. (5), we can proceed to compute the forces acting on each cell centre based on the VM energy functional in Eq. (1). This is done directly by computing the negative gradient of the energy. Therefore, the force on the centre of cell *i* is computed as

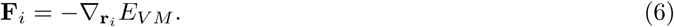

The expression for the gradient is made somewhat complicated by the fact that the derivative is taken with respect to the position of the cell centre, while the energy of the VM is naturally written in terms of the positions of the dual vertices. After a lengthy but straightforward calculation (see Appendix I) we obtain:

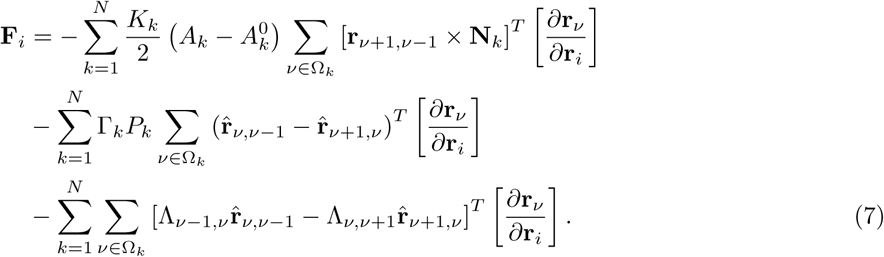

In the last expression, 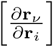 is the 3 × 3 Jacobian matrix (see Appendix I) connecting coordinates of cell centres with coordinates of the dual Voronoi tessellation and [·]*^T^*[·] denotes a row-matrix product producing a 3 × 1 column vector. Note that this product does not commute, i.e., the order in which terms appear in the expression above is important. *N* is the total number of cells in the system.

We note that most terms in the *k* sum in Eq. (7) are equal to zero, since each vertex coordinate **r***_ν_* depends only the cell centres **r***_k_* associated to its Delaunay triangle (see Fig. 2b). In other words, we only need to consider cell *i* and its immediate neighbours. For clarity, we outline the algorithm for computing the area term and note that perimeter and junction terms can be treated in a similar fashion.

1. For particle *i* compute 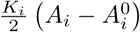 and multiply it by the sum 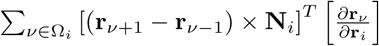. This sum is over all vertices (corners) *ν* of cell *i*.
2. For all immediate neighbours *j* of cell *i* compute 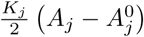 and multiply it by the sum

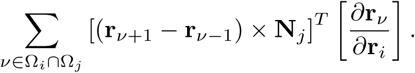

Note that *ν* ∈ Ω*_i_* ∩ ∈ Ω*_j_* ensures that vertices *ν* surrounding *j* are taken into account only if they are affected by (i.e., also belong to) the cell *i*.

From the previous discussion it is clear that the expression for the force is local, i.e., computing the force does not require including cell centre positions beyond the immediate neighbourhood of a given cell. This is extremely beneficial from a computational point of view as one can readily utilise standard force cutoff techniques, such as cell and neighbour lists^64^ in order to speed up force computations. It is also evident that the force in Eq. (7) is not pairwise, i.e., it cannot be written as a sum of forces acting on a pair of cells, or in mathematical terms, 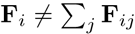. This is not surprising since the position of the Voronoi vertices depends on the position of three cell centres.

### 2. Cell alignment

The AVM also includes the *cell polarity*^65^ modelled as a unit length vector, **n***_i_*, laying in the *xy* plane. This vector determines the direction of the cell’s motion and division and should not be confused with apical-basal polarity, which is not included in this model. In the simplest case, the vector **n***_i_* does not take any preferred direction, but instead randomly fluctuates under the influence of uncorrelated random noise originating from the intercellular processes and the environment. This simple model can be augmented to include several different models for cell-cell alignment. For example, a cell can be given a tendency to align its polarity to its immediate neighbours by minimising the alignment energy,

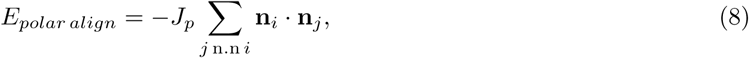

where *J_p_* > 0 is the alignment strength and the sum is over all nearest neighbours of *i*. This model is similar to the Vicsek model, which is used in studies of flocking, e.g. of swarms of birds.^66^The molecular details of polar alignment are not yet understood in detail. It as been argued that in some tissue systems this involves components of the planar cell polarity signalling cascade,^67,68^ but even here their precise coordination and involvement are as yet not resolved.

Alternative alignment mechanisms consider only internal processes in the cell and do not directly depend on the polarity of the neighbouring cells. One such mechanism introduced in physics of active matter^69,70^ assumes that cell polarisation vector aligns with the migration direction of the cell, i.e.,

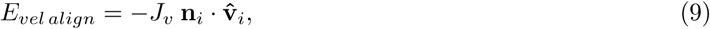

where **v̂***_i_* is the normalised cell velocity vector and *J_v_* > 0 is the alignment strength.

Finally, **n***_i_* can also align to the direction of the eigenvector of the cell shape tensor, i.e. the cell seeks align direction of **n***_i_* along its long axis. This is achieved by minimising the value of the energy

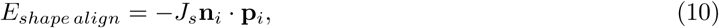

where *J_s_* > 0 is the alignment strength and **p***_i_* is the eigenvector of the matrix^71^

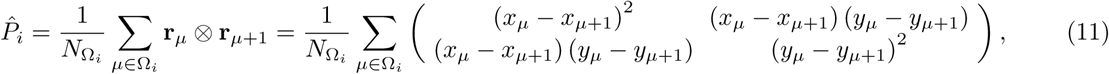

corresponding to its largest eigenvalue. This mechanism is one possible pathway to obtain *plithotaxis.* Each of these alignment mechanisms causes a torque on **n***_i_*, given by

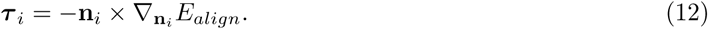

In the *SAMoS* implementation of the AVM, one can select which of these alignment mechanisms are to be included in an actual simulation.

### 3. Equations of motion

We proceed to write equations of motion for the position and orientation of cell *i* in the AVM. As discussed in Sec. II A, at the cellular scale inertial effects are negligible. In this overdamped limit, the equation of motion for the position of the cell centre is just force balance between friction and driving forces. If we assume that the friction of the cell with its surrounding and the substrate is isotropic and can be modelled by the single friction coefficient *γ*, the equation of motion for the position of cell centre *i* becomes

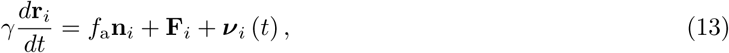

where **F***_i_* is the sum of all forces acting on the cell, i.e., a sum of the VM model forces given in Eq. (7) and the soft-core repulsion defined in Eq. (56) in Appendix I. ***ν**_i_*(*t*) is an uncorrelated stochastic force that models the effects of random fluctuations. These fluctuations originate from the intracellular processes and interactions with the environment. In its simplest possible form, one can assume that ***ν**_i_*(*t*) has no time correlations, i.e., each cell experiences a random “kick” at a given instant of time whose magnitude and direction do not depend on any past “kicks” on that cell. Formally, we write 〈*ν_i,α_*(*t*)〉 = 0 and 〈*ν_i,α_*(*t*) *ν_j_*,*β* (*t*′)〉 = 2*Dδ_ij_δ_αβ_δ* (*t* – *t*′), where *D* measures the average magnitude of the stochastic force and is interpreted as an effective translational diffusion coefficient, *α, β* ∈ {*x,y*} and 〈… 〉 denotes the statistical average. Finally, the term *f*_a_**n***_i_* has its origins in the physics of active matter systems.^58^It models *activity*, i.e., internal cellular processes that drive it to move in the direction of its polarity. The active force strength *f*_a_ controls the magnitude of this activity. While the biological meaning of *f*_a_ may appear unclear, it quantifies a cell’s ability to move on its own due to the complex molecular machinery within it. In many models of individual cells crawling on a substrate with a prominent lamellipodium, the resultant active velocity *υ*_0_ = *f*_a_/*γ* is due either actin tread-milling,^72^ differential friction or differential contractility. In an epithelium, on the other hand, *f*_a_ should be understood as an effective parameter that models the non-balanced active force due to junction contractions and internal cell contractility.^73^ Although the molecular origin of *f*_a_ is at present not fully understood, even this simple description of the activity combined with the cell polarity alignment models discussed in the previous subsection leads to collective behaviour that resembles behaviour observed in experiments.

We then write equations for motion for the polarity vector. If we define the angle of vector **n***_i_* with the *x*-axis as *θ_i_*,then **n***_i_* = (cos *θ_i_*, sin *θ_i_*)and we have

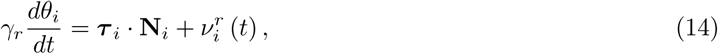

where ***τ**_i_* is the torque acting on **n***_i_*, given in Eq. (12), **N***_i_* is the local normal to the cell surface (i.e., simply the unit length vector in the *z*-direction), *γ_r_* is the orientational friction, and 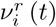 is responsible for introducing an orientational randomness. Akin to the stochastic force ***ν**_i_*(*t*) in Eq. (13), 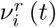 (t) has properties 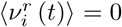 and 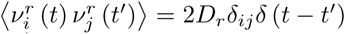, where *D_r_* is an rotational diffusion constant. The related time scale *τ_r_* = *γ_r_*/2*D* measures the time necessary for the cell polarisation direction to decorrelate due to fluctuations produced by cellular processes and the environment. The unit-length vector **n***_i_* then evolves according to 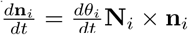.^74^

Eqs. (13) and (14) describe the cell dynamics in the AVM. These equations are solved numerically using several standard time discretisation schemes.

### 4. Cell growth, division and death

Another contribution to the activity comes from cell growth, division and death or extrusion from the cell sheet, present in many epithelial tissues. We will refer to these three processes as *population control*. Cell growth is modelled as a constant-rate increase of the native area, i.e.,

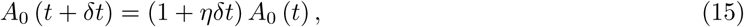

where *η* is the growth rate per unit time and *δt* is the simulation time step. Cell death is modelled by tracking the cell’s “age”. Age is is increased at every time step by *δt.* Once the age reaches a critical value, the cell is removed from the system. We note that onset of the cell’s death (or alternative its extrusion from the sheet) may also depend on the stresses or forces exerted on the cell. In the current implementation of the AVM we do not include such effects. Adding such effects to the model would be straightforward.

Finally, cell division is modelled based on the ideas of Bell and Anderson.^75^ The cell can only divide once its area is larger then some critical area *A_c_*, in which case it divides with probability proportional to *A* – *A_c_*,
i.e.,

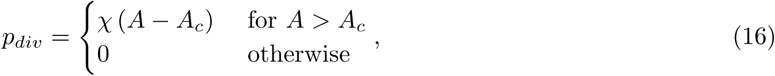

where *χ* is a constant with units of inverse area times time. Upon division the two daughter cells are placed along the direction of the vector **n***_i_* and their ages are reset to zero. The Delaunay triangulation is rebuilt after every division or death event.

The cell cycle is clearly a far more complex phenomenon than what is captured by this simple model. The simplest possible extension would be to build upon the ideas of Smith and Martin,^76^ or use many other more sophisticated models available in the literature. Adding such extensions to the AVM would be straightforward and will be included in later versions of the model.

### 5. Maintaining a Delaunay triangulation/Voronoi tessellation

In an actual simulation, we start either from carefully constructed initial positions of cell centres or choose cell positions from a particular experimental system. Those positions are then used to build the initial Delaunay triangulation and its corresponding dual Voronoi tessellation. However, as the cell centres move, there is no guarantee that the Delaunay character of the triangulation is preserved. For the model to be able to properly capture cell dynamics we, however, need to ensure that the triangulation is indeed Delaunay at each time step. Building a new triangulation in every time step would be computationally costly. Instead, we apply the so-called equiangulation procedure:^77^ For every edge in the triangulation we compute the sum of the angles opposite to it. If the sum is larger than 180° we “flip” the edge (see Fig. 2c). The procedure is repeated until there are no more edges left to flip. One can show^77^ that this procedure always converges and leads to a Delaunay triangulation. While equiangulation is not the most efficient way to build a Delaunay triangulation “from scratch”, if one starts with a triangulation that is nearly Delaunay, in practice only a handful of flip moves are required to recover the Delaunay triangulation. Given that cell centres move continuously, in the AVM the equiangulation procedure significantly increases the efficiency of maintaining the Delaunay triangulation throughout the simulation.

It is important to note that edge flips are local and flipping one edge in the Delaunay triangulation only affects one edge of the dual Voronoi tessellation. This means that a single edge flip can only affect junctions between four cells. The centres of those four cells coincide with the locations of the four corners of the polygon formed by two triangles sharing the flipped edge (see, Fig. 2c). This is precisely the mechanism behind the T1 transition discussed in Sec. IIIA below. No other cell contacts are affected by the flip.

### 6. Handling boundaries

The implementation of the SPV by Bi,*et al.*^57^assumes periodic boundary conditions. This assumption is reasonable if one studies a relatively small region of a much larger epithelial tissue. Many experiments, especially those of cell migration on elastic substrates, however, are performed with a relatively small number of cells where the effects of boundaries cannot be neglected or are even the main focus of the study. Therefore, in the AVM we include an open, flexible boundary.

In order to avoid very costly checks of topology changes, we assume that the topology (connectivity) of the boundary is maintained throughout the entire simulation. This does not mean that the boundary is fixed. It can grow, shrink and change its shape, but it cannot change its topology, e.g. it is not possible to transition from a disk to an annulus. Examples of allowed and disallowed changes of the boundary are shown in Fig. 3. While fixing the boundary topology may appear restrictive, we will see below that in practice a model with fixed topology of the boundary allows for detailed studies of a broad range of problems directly applicable to many current experiments.

**Figure 3.**
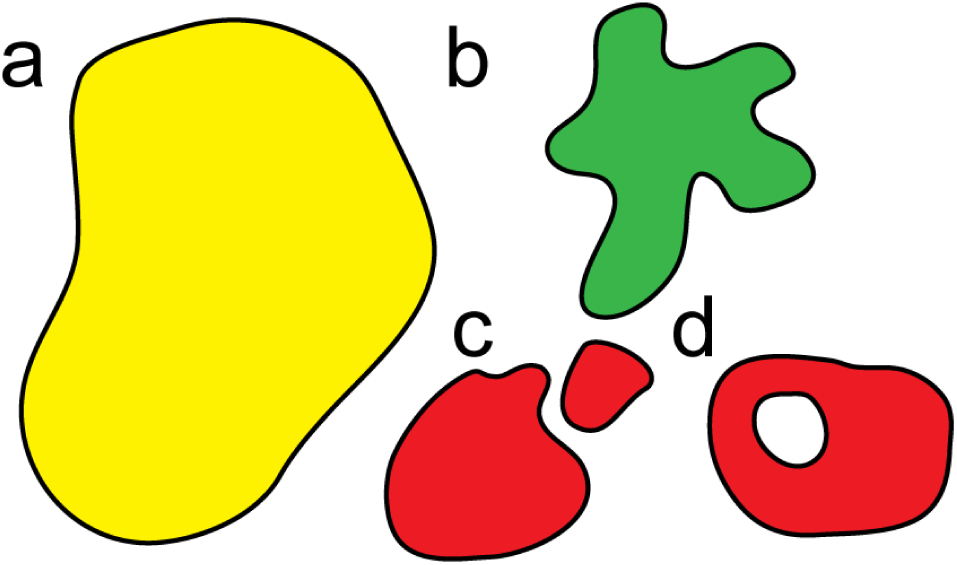
Schematic representation of the allowed and disallowed changes of the boundary in the AVM. An initial configuration with the topology of a disk (a) is allowed to develop pronounced fingers (b). However, it is not possible to split into two domains (c) or develop a hole (d), which would both lead to the introduction of a new boundary lines, and therefore lead to changes in the topology.

In the AVM, the degrees of freedom are cell centres, represented as particles. These particles serve as sites of the Delaunay triangulation. In order to handle boundaries we introduce a special type of *boundary* particle and in addition, we also specify the connectivity between the boundary particles. Boundary particles together with their connectivity information form a *boundary line*. This line sets the topology of the boundary, which is preserved throughout the simulation, and delineates between the tissue and its surrounding. In addition, the boundary line can have an energy associated to it. Biologically, this boundary energy corresponds to the complex molecular machinery, such as actin cables, that is known to affect the behaviour of the free edge of an epithelial sheet. Specifically, we introduce the boundary line tension,

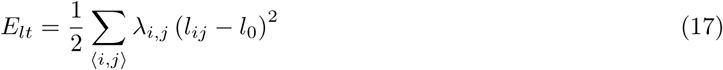

where λ*_i_,_j_* is the line tension of the edge connecting boundary vertices *i* and *j*, *l_ij_* = |**r***_i_* – **r***_j_*| is the length of that edge and *l*_0_ is its native (preferred) length. In addition, we also introduce the boundary bending stiffness

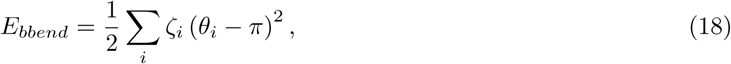

where *ζ_i_* is the stiffness of angle *θ_i_* at the boundary particle *i*, determined as

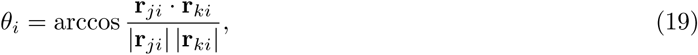

where **r***_j_* and **r***_k_* are the positions of boundary particles to the left and to the right of particle **r***_i_*. For simplicity, we assume that each boundary particle has exactly two boundary neighbours. This prevents somewhat pathological, “cross-like” configurations where two otherwise disjoint domains would hinge on a single boundary site.

It is important to note that boundary particles do not represent centres of an actual cell. The Voronoi polygon dual to a boundary particle in a Delaunay triangulation extends out to infinity. Consequently, it is not possible to unambiguously assign quantities such as the associated area or perimeter to boundary particles. Therefore, only internal particles correspond to cell centres, while boundary particles should be thought of as “ghosts” that serve to mark the edge of the cell sheet. These particles, however, experience forces from the interior of the tissue as well as from the interactions with their neighbours on the boundary. The boundary line is able to dynamically adjust its shape and length in this manner. We also allow for boundary particles to be added to or removed from the boundary line. Details of the algorithms used to dynamically update the boundary are given in Appendix II.

This concludes the description of the AVM. In Fig. 4 we present the flow chart of the main steps in computing the time evolution of cell centres and polarity vectors. Technical details of the implementation are given in Appendix III and in the on-line documentation provided with the *SAMoS* code.

**Figure 4.**
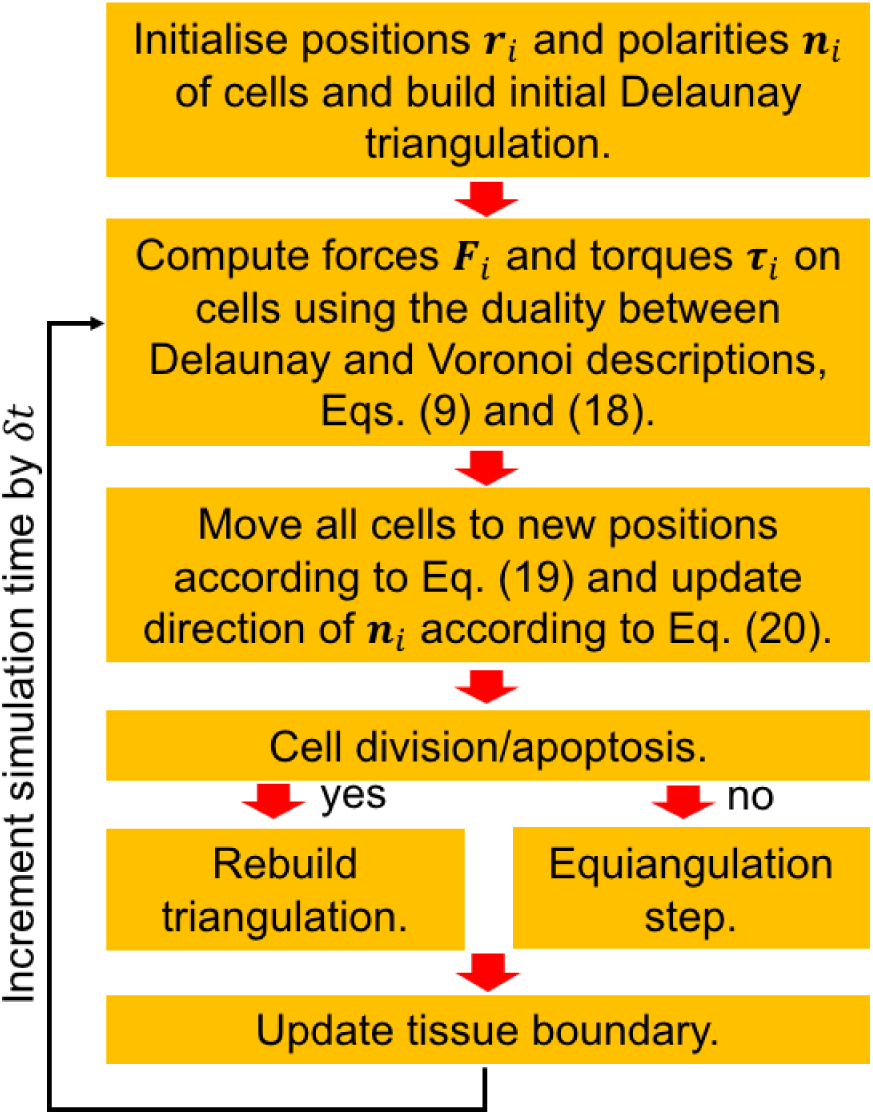
Flow chart of the force and torque calculations in each time step based on Eqs. (7) and (12). The calculation speed is significantly improved by using the fast equiangulation moves shown in Fig. 2c. The Delaunay triangulation is recomputed only if there are cell division and/or death events, which do not occur at every time step.

## III. EXAMPLES AND APPLICATIONS

In order to validate the model and compare it with the results of similar models proposed in the literature, as well as to show its use in modelling actual biological tissues, we now apply the AVM to several problems relevant to the mechanics of epithelial tissue layers.

### A. T1 transitions

We start by illustrating one of the key processes observed in epithelial tissues, the T1 transition. As detailed in Fig. 2, in the AVM T1 transitions are handled through an edge flip in the Delaunay triangulation. An edge flip only happens when, in the notation of Fig. 2c, we have *α* + *β* = *δ* + *γ* = 180°. Then both triangles are circumscribed by the same circle passing through its combined four vertices. The location of the T1 transition coincides with the centre of this circle. Due to the continuous connection between the position of sites of the Delaunay triangulation and its dual Voronoi tessellation, we always approach this point smoothly, i.e. a junction between two cells will smoothly shrink to a point, the T1 transition will occur, and then it will expand in a new direction. This process arises naturally in the AVM model, in stark contrast with many currently available implementations^78^ that require a cut-off criterion on the edge length of a cell before a T1 transition can occur. It also avoids discontinuous jumps at finite edge length, bypassing the T1 point altogether, also a feature of a number of models, notably those based on sequential energy minimisation.^47,79,80^ Next to its high computational efficiency, the ability to smoothly go through a T1 transition without the need for any additional manipulations of either the Delaunay triangulation or the Voronoi tessellation is one of the key advantages of the AVM approach.

In Fig. 5, we illustrate a T1 transition in the bulk, in a region of phase space where the system exhibits liquid-like behaviour, but with very slow dynamics (see next section). The edge linking cells 2 and 4 (in red) slowly shrinks to a point, and then rapidly expands in the opposite direction. This feature points to dynamics akin to certain models of sheared materials,^81^ where the active driving pulls the material over an energy barrier from one minimum to the next. It is somewhat different from the activated dynamics which has been proposed for the SPV,^79^ which would predict a series of fluctuations through which the barrier between minima is ultimately crossed.

**Figure 5.**
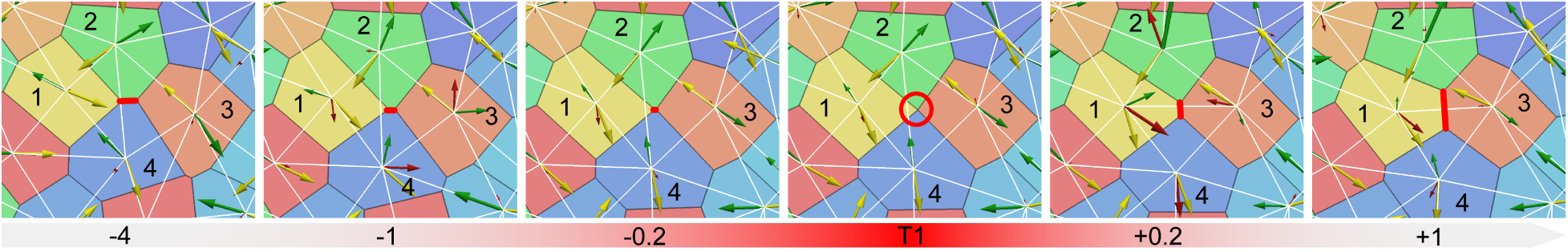
Time lapse of a T1 transition, each label is the time (in units of *γ*/*Ka*^2^, with *a* =1) with respect to the T1 event. Initially, cells 2 and 4 are in contact (red line). Approaching the transition, the connecting line slowly contracts, until it becomes a point at the transition. Cells 1 and 3 make a new contact which then rapidly expands. The arrows represent the force on the cell centre resulting from the Vertex potential (green), which partially compensates the active self-propulsion force (yellow) to give the resultant total force (red). The Voronoi tessellation is outlined in black, and the Delaunay triangulation is in white.

### B. Activity driven fluidisation phase diagram

We now explore different modes of collective behaviour, i.e., *phases*, of the tissue based on the values of parameters of the original VM (*K_i_*, Γ*_i_* and Λ*_μ_*,*_ν_*), and AVM-specific parameters such as the activity *f_a_*, the orientational correlation time *τ_r_*, and the boundary line tension λ. In order to keep the number of independent parameters to a minimum, it is again convenient to rewrite the energy of the VM, Eq. (1), in a scaled form.^47,57^ We first choose *K_i_* = 1 and set 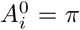 as an area scale. For simplicity, we assume that all perimeter and junction tensions are the same, i.e., we set Γ*_i_* ≡ Γ and Λ*_μ_*,*_ν_* ≡ Λ for all *i*, *μ* and *ν*. Then, as discussed below Eq. (1), we can complete the square on the second and third terms in Eq. (1) and obtain the scaled VM potential

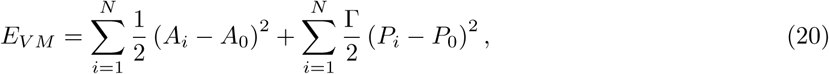

where *P*_0_ = –Λ/Γ and *A*_0_= *π*. The first term in Eq. (20) penalises changes in the cell area, while the second term penalises changes of the perimeter. There is no reason for the preferred area *A*_0_ to be generically compatible with the preferred perimeter *P*_0_. This sets up a competition between the two terms in Eq. (20), giving a natural scale that is is determined by the relative ratio of 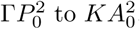. In other words, if 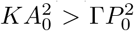, the cell will try to optimise its area at the expense of paying a penalty for not having the most optimal perimeter, and the opposite if 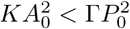.

Bi, *et al.^57^* introduced the dimensionless *shape factor* 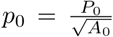, which controls the ratio of the cell’s perimeter to its area through the target area *A*_0_ and target perimeter *P*_0_. The value of *p*_0_ then determines whether the area or the perimeter term in Eq. (20) wins and effectively sets the preferred shape of each cell: cells of different shapes have different values of *p*_0_. For example, regular hexagons, pentagons, squares and triangles corresponds to *p*_0_ = 3.722, *p*_0_ = 3.812, *p*_0_ = 4.0 and *p*_0_ = 4.559, respectively.

Remarkably, one observes^47,57,79^ a transition between a solid-like behaviour of the tissue, where cells do not exchange neighbours, and liquid-like behaviour, where neighbour exchanges do occur, at *p*_0_ = 3.812, a value that corresponds to a regular pentagon. At present, the biological significance of this observation is not clear, but it appears to be a robust feature of many experimental systems.^82^In order make the comparison between the AVM and the SPV model, we also adopt *p*_0_ as a main parameter that controls the preferred cell shape.

In order to initialise the simulation, in each run, we start by placing soft spheres with slightly polydisperse radii in a circular region. We then use *SAMoS* to minimise the energy of a soft sphere packing in the presence of a fixed boundary. This ensures that initially, cells are evenly spaced without being on a grid. We also fix the packing fraction to *ϕ* = 1, ensuring that the average cell area of the initial configuration is 〈*A*〉 = *A*_0_. The boundary is either fixed (referred to as “fixed system”), or allowed to fluctuate freely (“open system”). Fig. 6 shows a representative set example of the states that we observe. We run the simulation for either 100,000 time steps with step size *δt* = 0.01 in the unstable region (e.g., Fig. 6e), or 250, 000 time steps with *δt* = 0.025 in the solid-like region (e.g., Fig. 6c). For these systems with *N* = 1000 cells in the interior, this takes between 10-40 minutes on a single core of a modern Intel Xeon processor depending on the number of rebuilds of the Delaunay triangulation that are necessary (more in the liquid-like phase).

**Figure 6.**
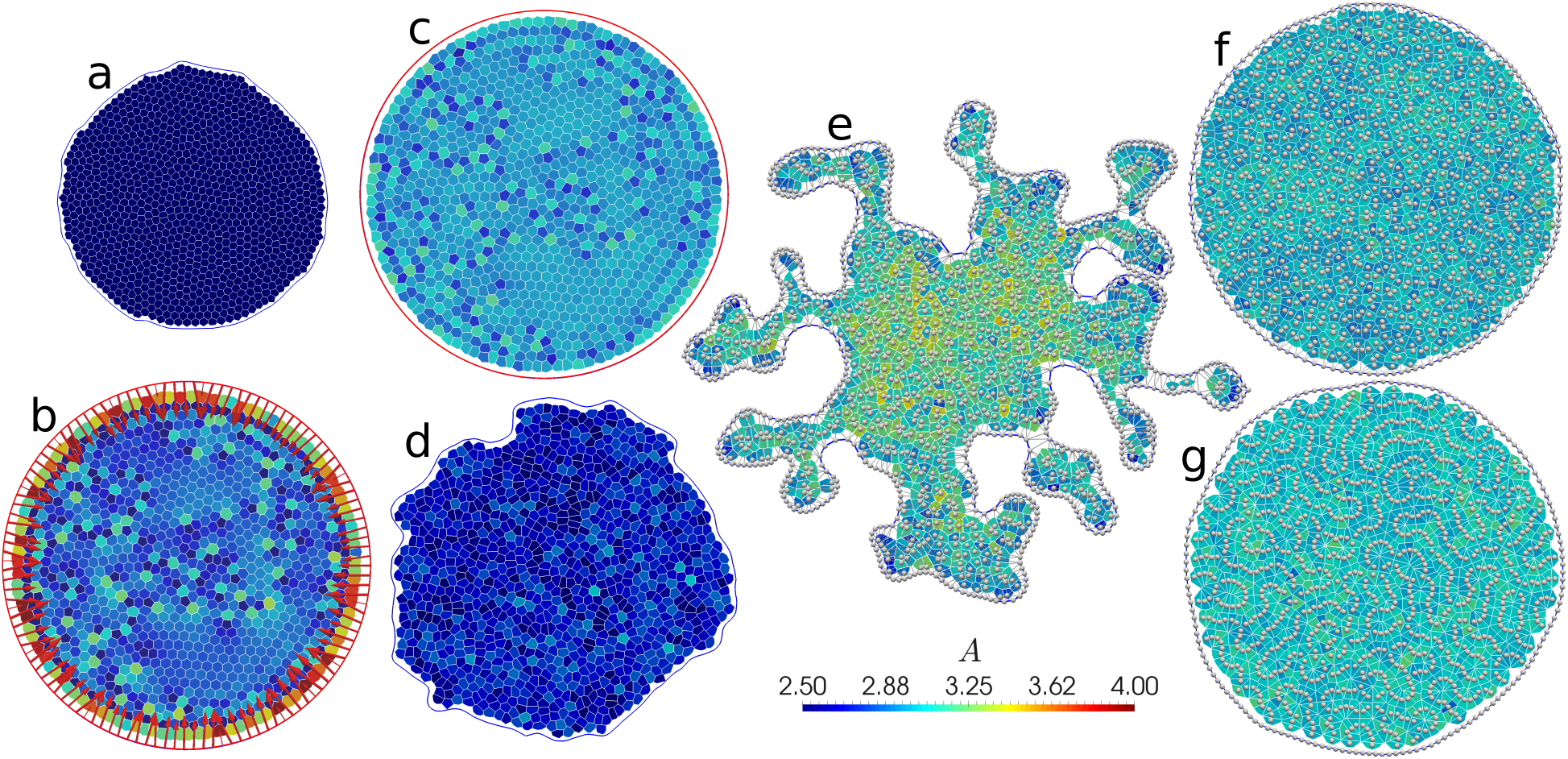
Representative snapshots of states seen with fixed and flexible boundaries in the AVM. Contact lines between cells are outlined in white, and cells are coloured according to their area. The line connecting tissue boundary points is blue for flexible boundaries (panels a, d, e, f and g), and red for fixed ones (panels b and c). In panels e, f and g, cell centres are denoted by white spheres, and in panels e and f, the Delaunay triangulation is also shown as grey lines. (a) Shrinking cells at *p*_0_ = 2.48, Γ = 1.0, 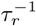 =0.01 and *f*_a_ = 0.1, no boundary line tension. (b) Same as (a),but for a fixed boundary. (c) Solid-like (glassy) state at *p*_0_ = 3.39, Γ = 1.0, 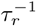 =0.1 and *f*_a_ = 0.03. (d) Liquid-like state with a fluctuating boundary at *p*_0_ = 3.39, Γ = 1.0, 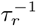 = 0.1 and *f*_a_= 0.3, no boundary line tension (e) Fingering instability at *p*_0_ = 3.72, Γ = 0.1, 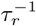 = 0.01 and *f*_a_ = 0.3, boundary line tension λ = 0.1. (f) Fluid state at *p*_0_ = 3.95, Γ = 0.1, 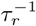 = 0.1 and *f*_a_= 0.1, boundary line tension λ = 0.3. (g) Rosette formation at *p*_0_ = 4.85, Γ = 0.1, 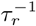 =0.1 and *f*_a_ = 0.03, boundary line tension λ = 0.3.

The unit of time is set by *γ*/*K_a_*^2^, where *a* ≡ 1 is the unit of length. We note that Bi, *et al.* use 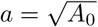 as the unit of length. This is possible as long as cells are not allowed to grow, i.e., when *A*_0_ changes in time. The AVM allows for the cells to change their size and therefore we need to choose a different unit of length. In our case, *a* is the range of the of soft-core repulsion between cell centres (see, Eq. (56) in Appendix I).

At low values of *p*_0_, we find a system that prefers to be in a state with mostly hexagonal cells, unless the active driving *f*_a_ is very high. Open systems will shrink at this point so that all cells are close to their target *P*_0_, as shown in Fig. 6a. Consistent with this, larger values of the perimeter modulus Γ lead to stronger shrinking. For fixed systems, this route is blocked, and instead there is a strong inward tension on the boundaries and a gradient in local density, as shown in Fig. 6b.

In agreement with the results of Bi, *et al.,*^57^we find that at low *p*_0_ < 3.81 and low values of driving *f*_a_, cells do not take an organised pattern and do not exchange neighbours. Recast in the language of solid state physics, the tissue is in an *amorphous solid* or *glassy* state. In Fig. 6c we show such a state for a fixed boundary. In order to characterise the physical properties of this state, we measure the dynamical time scale of cell rearrangements through a standard tool of the physics of glassy systems, the self-intermediate scattering function^15^

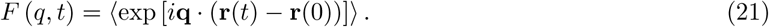

*F* (*q*, *t*) measures the decay of the autocorrelation of cell-centre positions **r** (*t*) at a particular wave vector, **q**, taken usually to be the inverse cell size *q* ≡ |**q**| = 2*π*/*a*. The long-time decay of *F* (*q*,*t*) is characterised by the so-called *alpha-relaxation time τ_α_* at which *F* (*q*, *t*) has decayed by half. When the system solidifies, i.e. when neighbour exchanges stop, **r** (*t*) remains constant and hence *τ_α_* diverges,^15^ and stays infinite within the solid phase. In Fig. 7a-c, we show the phase diagram of *τ_α_* as a function of *p*_0_ and *f*_a_, for several systems with different boundary conditions.

**Figure 7.**
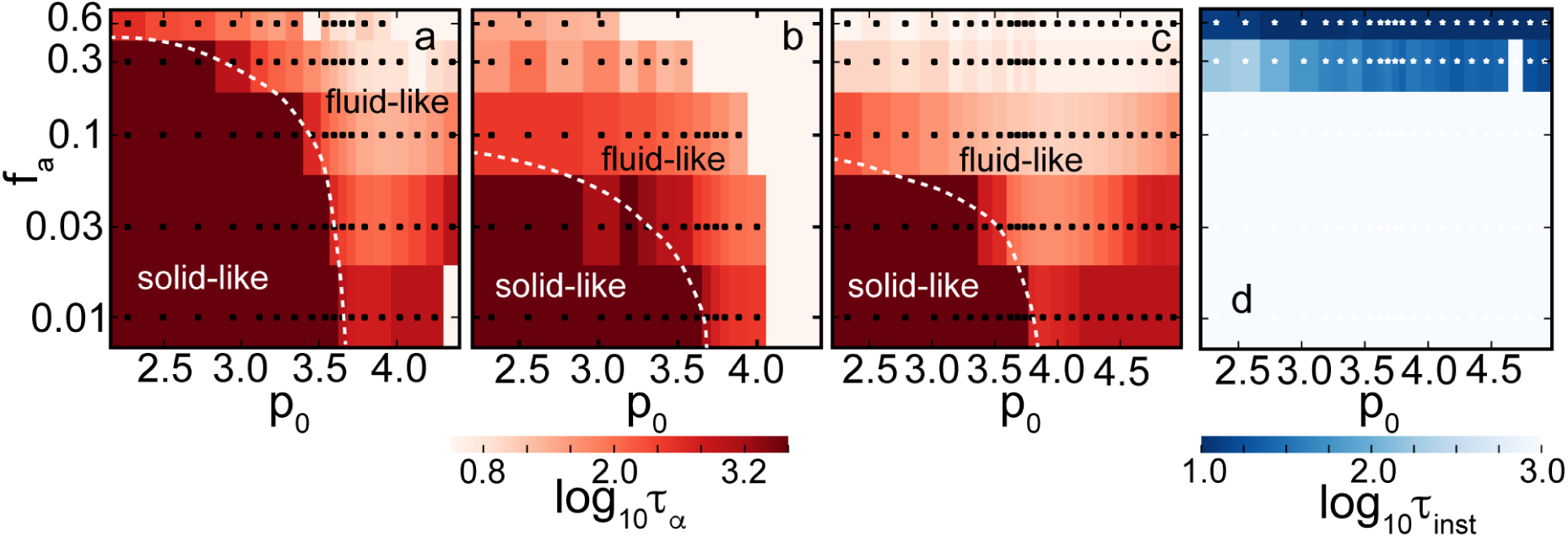
Phase diagrams of the AVM. Panels (a), (b) and (c): *α*—relaxation time *τ_α_* determined from the selfintermediate scattering function, Eq. (21). These plots indicate that it is possible to initiate cell intercalation events by changing values of *p*_0_ or active driving. High values of *τ_α_* (dark regions) correspond to the solid-like (glassy) phase where T1 events are suppressed. (a) Fixed system with Γ = 1 and 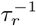 = 0.01. (b) Open system with Γ = 1, 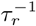 = 0.01 and boundary line tension λ = 0. (c) Open system with Γ = 0.1, 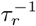 = 0.1 and boundary line tension λ = 0.1. All systems are solid-like at low *p*_0_ and low driving *f*_a_. The critical *p*_0_ = 3.81 is the same in (a), (b) and (c), but the open system becomes liquid-like at much lower values of active driving. Lowering Γ also lowers this transition point. The dashed white line represents a rough boundary between solid-like and fluid-like behaviour.(d) Characteristic time scale *τ*_inst_ needed to reach the boundary instability, determined from reaching a threshold boundary length (see text), for the same parameters as (c). Sufficiently high driving always leads to an unstable system.

In Fig. 7a we show regions of solid-like and liquid-like phases in a system with fixed boundaries, at Γ = 1, and a low noise value of *τ_r_* = 0.01. We find a boundary of the solid-like phase that stretches from *p*_0_ ≈ 3.81 at small *f*_a_ to a maximum activity *f*_a_ beyond which the system is fluid at all *p*_0_. This is qualitatively, but not quantitatively consistent with the results of Bi, *et al.,* who find a transition line at roughly twice our *f*_a_ values. Several factors are likely implicated in this discrepancy. Our systems, at *N* = 1000 cells are more than twice as large as the *N* = 400 systems considered by Bi, et *al.,* and finite system size effects seem to play an important role, as shown below. We measure *τ_α_* at a value of 1/*q* corresponding to displacements of one cell size. However, even though displacements are large, we have evidence that this may not be sufficient to induce T1 transitions and therefore fluidise the system. Finally, fixed boundaries were used here and the periodic boundaries of Bi, et *al.* are likely not strictly equivalent.

The influence of the type of boundary conditions is very significant. In Fig. 7b, we show the phase diagram for the same Γ = 1 and 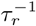 = 0.01 as in Fig. 7a, except with open boundary conditions and boundary line tension λ = 0. Separately, for Γ = 0.1, we have also confirmed that the value of the boundary line tension does not significantly affect the onset of the solid-like regime (not shown). We find a significantly lower maximum *f_a_* for the transition, *f*_a_ = 0.03, a factor of 10 compared to the fixed case. The effect also persists at 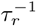 = 0.1, but is less pronounced (not shown). While we do not have a full explanation for this result, we do note that fluctuations of the boundary allow for rearrangements that are otherwise strongly suppressed by the fixed boundary. For example, the system in Fig. 6d shows significant boundary fluctuations. It is liquid-like with *τ_α_* ≈10, whereas the equivalent fixed system has *τ_α_* ≈ 100. In view of the significant role of the boundary, we expect a strong system-size dependence.^83^

At very high *p*_0_ and low active driving, we observe a systematic increase of *τ_α_* (especially visible in Fig. 7c). This unexpected result is accompanied by structural changes in the cell patterns that we observe. Fig. 6f shows a liquid system at *p*_0_ = 3.95, near the relative minimum *τ_α_* for a given *f*_a_. The distribution of cell centres appears random. In contrast, as can be seen in Fig. 6g, at very high *p*_0_ = 4.85, cells arrange themselves into rosette shapes, where many vertices meet in a point. Rosettes are a feature of many developmental systems,^50^ so it is interesting to see that they do appear naturally in the AVM context. Cell centres also arrange themselves in equidistant chains, hinting at a connection to one of the various pattern-formation instabilities studied in nonlinear dynamics. We note that this regime is numerically delicate, and the addition of the soft repulsive core between cell centres (see Appendix I) is necessary to make simulations stable. At present it is not clear if these effects are artefacts of the AVM, or have real biological significance.

In parts of the phase diagram, we observe a fingering instability^84 —87^ where regions a few cells wide migrate outward from the centre, as shown in Fig. 6e. When *τ_α_* drops below approximately 10, we observe that the fluctuations of the boundary already present in Fig. 6d become unconstrained. This is a mechanically unstable regime: Eventually, these cells will detach, a process we are not yet able to model due to the topological change that it would imply (see, Fig. 3). We have observed that fluctuations need to reach a threshold of approximately > 5% of a length increase in the boundary to break through to an unconstrained growth, otherwise the system remains stable, see e.g. Fig. 6d-g. We then associate a time scale *τ*_inst_ with reaching this threshold and use it to measure the degree of instability: a small time scale denotes a rapid growth rate of fluctuations. Fig. 7c shows *τ_α_,* and Fig. 7d shows *τ*_inst_ for the same open system with Γ = 0.1, 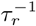 = 0.1 and boundary line tension λ = 0.1. We note that the transition line between solid-like and fluid-like states is low, at *f*_a_ = 0.03. At and below *f*_a_ = 0.1, the boundary of the system is stable, and above this threshold, the instability becomes more pronounced with increasing *f*_a_ and smaller *τ_α_*. The physical mechanism responsible for the instability involves a subtle interplay of *f*_a_, boundary line tension (stronger line tension suppresses the instability), the noise level 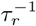 (lower 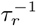 enhances the instability), and *p*_0_. The instability resembles observations of finger formation in MDCK monolayers.^84,85^ Existing models link it to either leader cells,^87,88^ a bending instability,^86^ or an active growth feedback loop,^89^ while here it emerges naturally. A detailed account of this phenomenon will be published elsewhere.^83^

### C. Growth and division

Division and death processes are important in any living tissue, for example, cell division and ingression processes play essential roles during development. Therefore, as noted in Sec. IIB 4, the AVM is equipped to handle such processes. It is important to note, however, that the removal of one cell during apoptosis or ingression and the addition of two new cells during division in the AVM causes a discrete change in the Voronoi tesselation which implies a discontinuous change of the local forces derived from the VM. We have simulated the growth of a small cluster of cells to assess whether this discrete change in geometry can lead to any instabilities in the model. These test runs did not reveal any artefacts due to discontinuities in the force caused by the division events.

In order to illustrate the growth process, we choose a shape factor, *p*_0_ = 3.10, corresponding to Γ = 1 and Λ = −5.5 and no active driving, i.e., we set *f*_a_ = 0. This puts the system into the solid-like phase where T1 events are absent. Our simulation runs for 10^6^ time steps at *δt* = 0.005, corresponding to 5000 time units, starting from 37 cells and stopping at about 24,000 cells. To balance computational efficiency with a smooth rate of division, cells are checked for division every 25 time steps. We show snapshots of different stages of the tissue growth in Fig. 8a-e.

**Figure 8.**
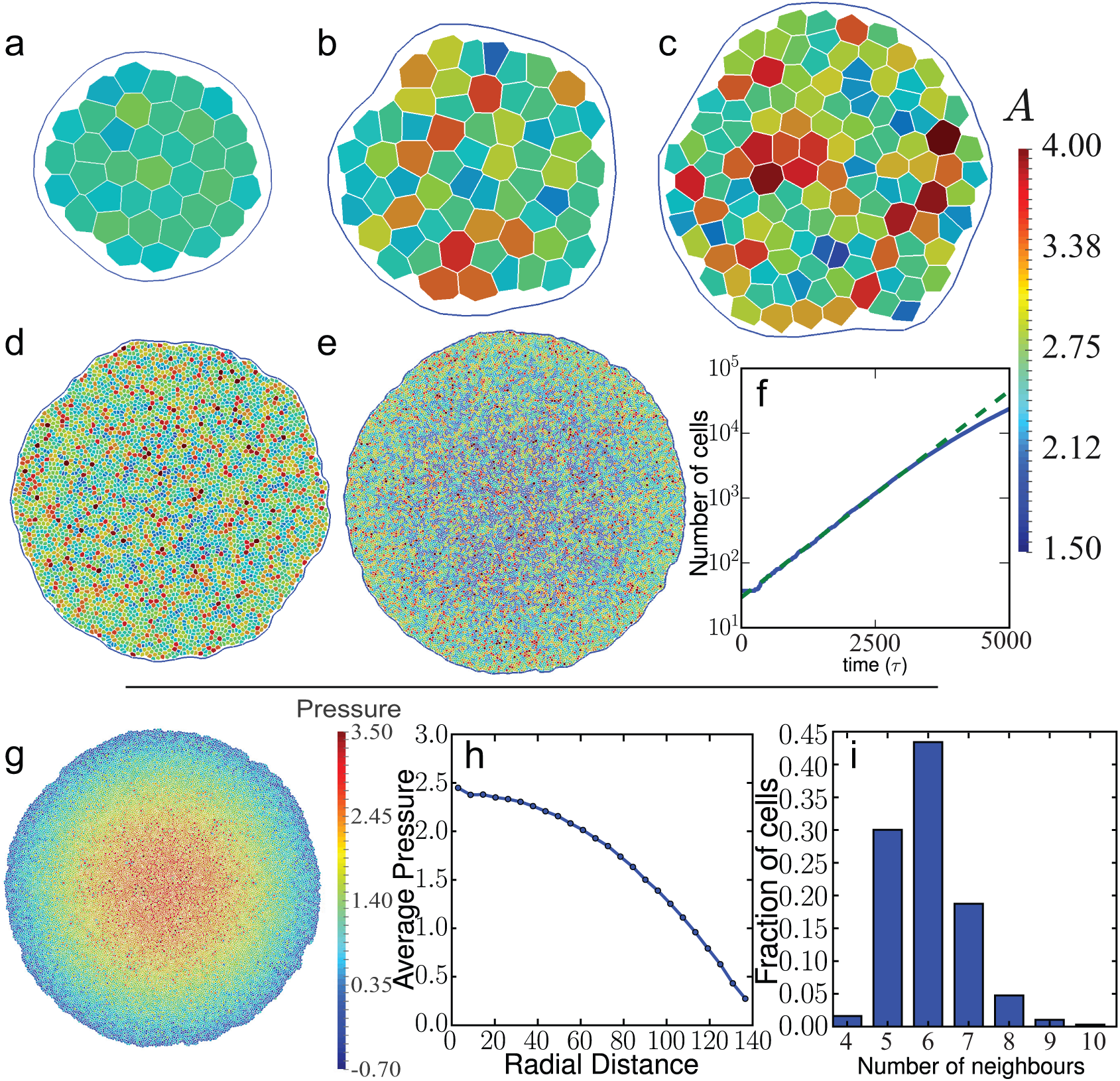
Snapshots of a growing epithelial tissue. Frames (a), (b), (c), (d) and (e) have 37, 63, 124, 4633 and 23787 cells and are at times 50, 500, 1000, 3500 and 5000, respectively. Cells have a chance to divide if their area is greater than a critical area *A_c_* = 2.8*a*^2^ after which the probability of a cell to divide increases linearly with its area (see Eq. (16)). In this simulation, the shape factor was set to *p*_0_ = 3.10. (f) Log-linear plot of the total number of cells as a function of simulation time. The growth rate of the patch is initially exponential but starts to slow at around 3000 cells. This is due to cells in the centre of the cluster being prevented from expanding by the surrounding tissue. (g) Tissue after 5000 time units (10^6^ time steps) with each cell coloured by pressure. Pressure has built up in the centre of the tissue while close to the edge the average pressure is low. (h) Average pressure (averaged over the polar angle) as a function of the radial distance from the centre of the tissue. (i) Distribution of the number of neighbours for cells in the system shown in (g).

We note that the numerical stability of the simulation that involves growth is quite sensitive to the values of the parameters used in the AVM. For example, divisions of highly irregularly shaped cells, as commonly observed in the high *p*_0_ regime, can put a significant strain on the simulation and even cause a crash. Helpfully, some of these problems can be alleviated with the help of the the soft repulsive potential defined in Eq. (56) in Appendix I between cell centres that acts to mediate the impact of cell divisions. Finally, as a rule, regardless of the exact parameter regime, a smaller time step is typically required for simulating growing systems.

In Fig. 8f we show the tissue size as a function of the simulation time. In this simulation there are no apoptosis or cell ingression events and, as expected, the tissue size grows exponentially. However, at long times, the growth slows down and deviates from exponential growth. This is easy to understand, as the centre of the tissue is prevented from expanding by the surrounding cells. The effect can be seen in Fig. 8e, where cells located towards the centre have, on average, smaller areas and in Fig. 8g, which shows a clear pressure buildup in the centre. This suggests that in the later stages, the simulated tissue is not in mechanical equilibrium any more. The pressure is computed using the Hardy stress description.^90^The details of the stress calculation in the AVM will be published elsewhere. We also see clear heterogeneities in the local pressure shown in Fig. 8g. In Fig. 8h, we show the radial pressure profile in the tissue at the end of the simulation. From the figure it is also evident that the there is a substantial pressure buildup close to the centre of the tissue as well as that angular averaging substantially reduces local pressure fluctuation notable in Fig. 8g. The origin of these effects warrants a detailed investigation and will be addressed in a later publication, we note however that stress inhomogeneities are a persistent feature of the epithelial cell monolayers that have been investigated by traction force microscopy.^12,91^

Finally, in Fig. 8i we show the distribution of the number of neighbours for this model system. The observed distribution is in a good agreement with the observations in actual tissues.^92,93^

### D. Modelling mechanically heterogeneous tissues

The AVM is equipped to allow for cell-specific parameters, which enables us to investigate tissues with locally varying mechanical properties. A commonly studied example of the effects such heterogeneities is cell sorting. As an example we show simulations that display sorting of two distinct cell types. We achieve this by setting the junction tension Λ for each pair of cell-cell and cell-boundary contacts. All our simulations consist of 1000 cells with half chosen randomly to be of the “red” type and the others being of the “blue” type. In these simulations, boundaries have been kept fixed. We observe sorting behaviour akin to that found in other commonly used tissue models.^35,94^ Using *r*, *b* and *M* to denote red, blue and the boundary, respectively, we start by fixing *K* = 1 and Γ = 1 and set −6.8 = Λ*_rr_* < Λ*_rb_* < Λ*_bb_* = −6.2, corresponding to p_0_ in the range 3.58 – 3.93. We, however, note that *p*_0_ is the quantity defined “per cell” and one should understand it in this context only as a rough estimate whether a given cell type is in the solid or fluid phase. All cells are subject to small random fluctuations of their position and polarity vector n*_i_*, which allows for T1 transitions that can bring initially distant cells into contact. We set the active driving to zero, i.e., *f*_a_ = 0. We have chosen different values for Λ*_rr_* and Λ*_bb_* to reflect the idea that the surfaces of these cells have different adhesive properties.^95^ Note that the Λ parameter for a particular contact is proportional to its energy per unit length. Sorting of cells into groups of the same type occurs when the energy of two red-blue contacts is greater than the energy of one red-red contact and one blue-blue contact, corresponding to Λ*_rb_* > (Λ*_rr_* + Λ *_bb_*) /2, see Fig. 9a-c. In this regime, for cells of the same type it is energetically favourable for the new contact to elongate while local red-blue contacts are shortened. Conversely, if Λ*_rb_* < (Λ*_rr_* + Λ*_bb_*) /2 then cells maximise their red-blue contacts forming a “checkerboard” pattern (Fig. 9d). The final pattern is not without defects, the number and location of which depend on the initial conditions. The tissue boundary consists of contacts between cell centres and boundary particles so Λ*_rM_* and Λ*_bM_* need also to be specified to reflect the way in which the cell types interact with the extracellular matrix or surrounding medium. Initially we set Λ*_rM_* = Λ*_bM_* = −6.2 and observe that blue cells cover the boundary enveloping red cells because this facilitates lower energy red-blue and red-red contacts being formed. If we incentivise red-boundary contacts by setting Λ*_rM_* < Λ*_rr_* + Λ*_bM_* −Λ*_rb_* = −6.6 we make red-boundary contacts preferable.^94^This case is shown in Fig. 9e for Λ*_rM_* = −6.8.

**Figure 9.**
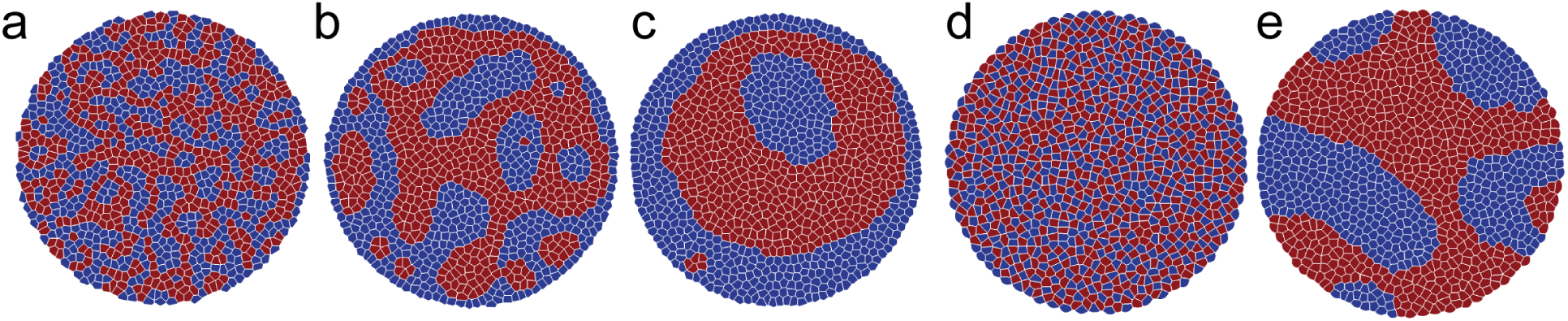
(a-c) Snapshots of a system of two cell types at times 10, 500 and 5000 with Λ*_rb_* = −6.4. (d) A “checkerboard” pattern formes immediately (at time 10) when red-blue cell-cell contacts are energetically favourable compared with pairs of red-red and blue-blue contacts, Λ*_rb_* = −6.7. (e) Same initial system as (a-c) but with red-boundary contacts slightly favoured over blue-boundary contacts. The system gradually separates into compartments of each cell type. The uncorrelated random fluctuations are sufficient to drive neighbour exchanges within the bulk of both the red and blue cell compartments. Cells on the compartment boundary can sometimes move parallel to it but meet strong resistance when trying to move across it.

### E. Effects of cellular alignment

We now briefly turn our attention to the effects of several models of cell polarity alignment. So far, we have assumed torque ***τ**_i_* = 0 in Eq. (14), i.e., we are in the situation where the the polarisation vector of each cell is independent of the surrounding cells and its direction diffuses randomly over time. In biological systems, it is known that a cell’s polarity responds to the surrounding and many forms of polarity alignment have been proposed. Here, we highlight two alignment mechanisms that are compatible with the current understanding of cell mechanics. In Fig. 10a-c, we have used the alignment model defined in Eq. (9) that assumes that the polarity vector **n***_i_* of cell *i* aligns with its velocity **v***_i_*. The torque term in Eq. (14) is then given by ***τ**_i_* = −*J_v_* **v***_i_* × **n**_i_, where *J_v_* is the alignment strength. This model was first developed for collectively migrating cells (modelled as particles),^69^ and it exhibits global polar migration, i.e. a state in which all particles align their velocities and travel as a flock. In dense systems of active particles confined to a finite region, velocity alignment has been shown to be intimately linked to collective elastic oscillations.^70^It is remarkable that the main hallmarks of this active matter dynamics are also observed in the model tissue. In Fig. 10a, we show velocity alignment dynamics for a confined system in the solid-like phase; here the collective oscillations are very apparent. They are strikingly reminiscent of the collective displacement modes observed in confined MDCK cell layers.^96,97^

**Figure 10.**
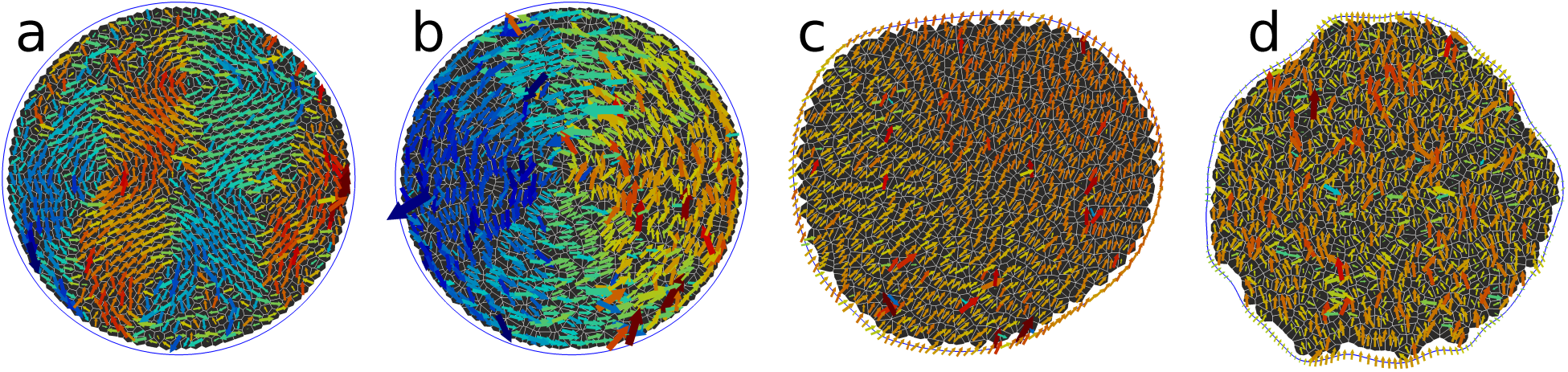
Effects of alignment. (a-c) are self-alignment of the cell polarisation with the cell velocity. (d) is self-alignment of the cell polarisation with the long axis of the cell. (a) A confined system in the solid-like region (*f*_a_ = 0.03, *p*_0_ = 3.385, 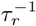 = 0.01), at alignment strength *J_v_* = 1.0 shows oscillating collective modes. (b) In the liquid region (*f*_a_ = 0.1, *p*_0_ = 4.4, 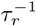 = 0.1), the system enters a vortex rotation state instead (*J_v_* = 1.0). (c) An open system at *f*_a_ = 0.1,*p*_0_ = 4.4, 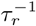 = 0.1, boundary line tension λ = 0.1 and *J_v_* =0.1 migrates collectively. (d) An open system at *f*_a_ = 0.1, *p*_0_ = 3.95, 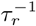 = 0.1, boundary line tension λ = 0.1 and alignment with cell shape with *J_s_* = 1 migrates collectively with complex fluctuations. Arrows are cell velocity vectors **v***_i_*; they are coloured by the magnitude of *υ_x_* for (a) and *υ_y_* for (b-d). Orange is positive (pointing right/upwards) and blue is negative (pointing left/downward).

In Fig. 10b, we apply the same dynamics, but now to a system that is in the liquid-like phase, with fixed boundaries. Here, the collective migration wins, but the confinement to a disk with fixed boundaries forces the cells into a vortex-shaped migration pattern. Finally, in Fig. 10c, in the absence of confinement, we recover the collective polar migration of the cell patch.

In Fig. 10d, we show the effects of aligning the cell’s polarity to the largest principal axis of the cell shape tensor, defined in Eq. (10). This type of alignment also leads to collective motion in an unconfined system, however there are significant fluctuations as the allowed cell patterns are highly frustrated by the constraint to remain in a Voronoi tesselation.

These preliminary results serve as a showcase of the non-trivial effects of cell-cell alignment on the collective behaviour of the entire tissue. A more detailed account of the effects of different alignment models will be published elsewhere.

### F. Modelling non-circular tissue shapes

In the previous discussions, all examples assumed a patch of cells with the topology of a circle. However, the AVM is not restricted to the circular geometry and can be applied to systems of arbitrary shapes, including domains with complex connectivity. Such situations often arise when modelling experimental systems where cells surround an obstacle, or in studies of wound healing. In Fig. 11 we present a gallery of non-circular shapes that can be readily studied using the AVM as it is implemented in *SAMoS.* The annular geometry shown in Fig. 11a would be suited for modelling wound healing problems as well as situations where cells migrate in order to fill a void. A common experimental setup where cell colonies are prepared as rectangular strips^84^ is shown in Fig. 11b, where three separate patches grow towards each other. Finally, in Fig. 11c we show an example of yet another very interesting situation, where cells are grown in a confined region of space.

**Figure 11.**
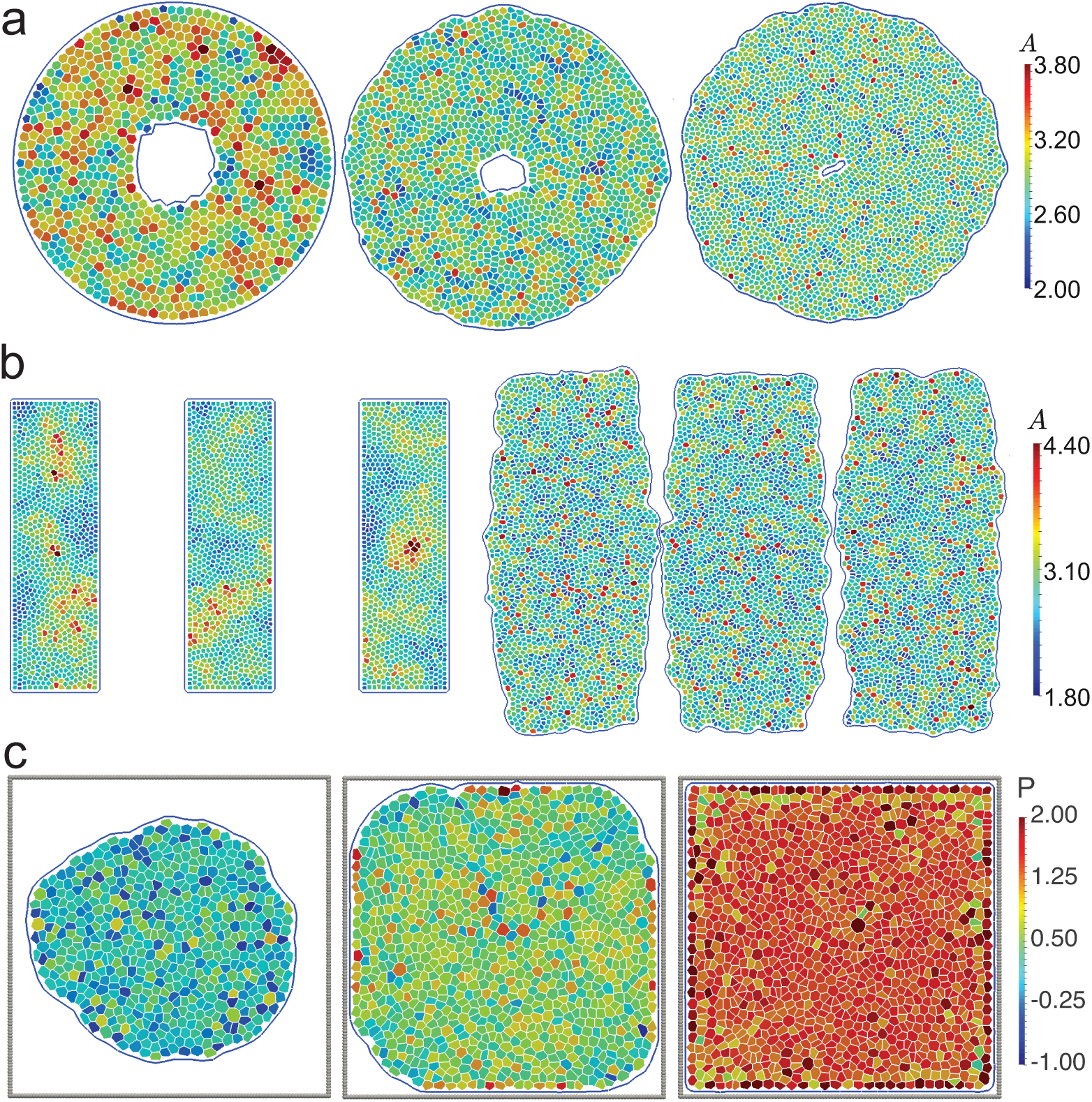
(a) Snapshots of the simulation of a cell sheet with an annular geometry used to illustrate how cells divide and fill the circular void in the centre. The system configuration was recorded at times 0, 1000 and 1700. The initial configuration has *p*_0_ = 3.46, i.e., it was in the solid-like phase. While it is not simple to define *p*_0_ for a growing system due to a constant change in the cell target area *A*_0_, we note that throughout the simulation, cell shapes remain regular. (b) Illustration of a common experimental setup where cells are grown in rectangular “moulds” for a system with initial *p*_0_ = 3.35. Once the entire region is filled with cells, the mould is removed and the colony is allowed to freely grow. Images on the right show two of the strips about to merge. In (c), we model tissue growth in confinement for a system with initial *p*_0_ = 3.46. Grey beads form the boundary of the confinement region that constrain the cell growth. Initially, cells do not touch the wall and freely grow. As the colony reaches the wall, one starts to notice pressure buildup. Finally, the entire cavity is filled with cells and any subsequent division leads to increase of the pressure in the system. Snapshots were recorded at times 1200, 1480 and 1600.

## IV. SUMMARY AND CONCLUSIONS

In this paper we have introduced the Active Vertex Model. It is a hybrid model that combines ideas from the physics of active matter with the Vertex Model, a widely used description for modelling confluent epithelial tissues. Active matter physics is a rapidly growing field of research in soft condensed matter physics, and it is emerging as a natural framework for describing many biophysical processes, in particular those that occur at mesoscales, i.e., at the scales that span multiple cells to the entire tissue. Our approach is complementary to the recently introduced Self-Propelled Voronoi model,^57^ for it allows modelling of systems with fixed and open, i.e. dynamically changing boundaries as well as cell-cell alignment, cell growth, division and death. The AVM has been implemented into the *SAMoS* software package and is publicly available under a non-restrictive open source license.

The AVM utilises a mathematical duality between Delaunay and Voronoi tessellation in order to relate forces on cell centres to the positions of the vertices of the dual lattice, i.e. meeting points of three of more cells - a natural description of a confluent epithelial tissue. This not only allows for a straightforward and efficient implementation using standard algorithms for particle-based simulations, but provides a natural framework for modelling topological changes in the tissue, such as intercalation and ingression events. In other words, in the AVM T1 transition events arise spontaneously and it is not necessary to perform any additional steps in order to ensure that cells are able to exchange neighbours.

Furthermore, our implementation of the AVM is very efficient, allowing for simulations of systems containing tens of thousands of cells on a single CPU core, thus enabling one to probe collective features, such as global cell flow patterns that span length-scales of several millimetres. In addition, the AVM is also able to handle multiple cell types and type specific cell contacts, which allows simulations of mechanically heterogeneous systems.

All these features make the AVM a strong candidate model to address many interesting biological and biophysical problems related to the mechanical response of epithelial tissues, especially those that occur at large length and time scales that are typically only accessible to continuum models. Unlike in the case of continuum models where relating parameters of the model to the experimental systems is often difficult and unclear, the AVM retains the cell-level resolution, making it simpler to connect it to the processes that occur at scales of single cells. The AVM is, however, not designed to replace continuum models, but to serve as the important layer that connects the complex molecular processes that occur at the cell level with the global collective behaviour observed at the level of the entire tissue.

With that in mind, there are, of course, still many ways the model can be improved. For examples, it would be very interesting to augment the AVM to include the effects of biochemical signalling. This would require solving a set of differential equations for signals in each time step, and then supplying those solutions to the mechanics part of the model. Adding such functionality would substantially increase the computational cost of simulations, however at the same time it would allow for detailed studies of the coupling between chemical and mechanical signalling. These are believed to be essential for developing a full understanding of the mechanics of epithelial tissues. Given the layout of the AVM and its implementation, implementing such functionality would be straightforward.

Furthermore, in the current version of the AVM activity is introduced in a very rudimentary manner, via assuming that cells self-propel in the direction of their polarity vector. This is a very strong assumption that would need a much stronger experimental support than currently available. It is also possible that a far more sophisticated model would be required to fully capture cell’s motility. However, one also needs to keep in mind that there is a tradeoff between being as biologically accurate as possible and retaining a sufficient level of simplicity to be able to efficiently perform simulations of large systems. With all this in mind, we argue that the our simulations clearly show that even this simplistic model of self-propulsion is capable of capturing many features of real systems, and that it can serve as a good starting point for building biologically more accurate descriptions.

Another potentially very interesting feature that is currently not supported would be splitting and merging of the boundaries, that is, allowing for topological changes of the entire sheet such as those depicted in Fig. 3c. This would us to study detachment of a part of the tissue or opening apertures as well as the opposite problem of closing holes and gaps. The latter is of great importance for studying problems related to wound healing. Unfortunately, setting up a set of general rules on how to automatically split a boundary line or merge two boundaries into a single one is not a simple task from a point of view of computational geometry. The problem is further complicated if those rules are also to be made biologically plausible, which is essential for the model to be relevant to actual experiments.

We conclude by noting that extending the model to a curved surface or making it fully three-dimensional would be more involved. Being able to study curved epithelial sheets, however, would be of great interest to systems where curvature clearly cannot be ignored, e.g., as in the case in modelling intestinal crypts.^98^ While there is nothing in the description of the AVM that is unique to the planar geometry, there are several technical challenges associated with directly porting it onto a curved surface. Most notably, building a Delaunay triangulation on an arbitrary curved surface is not a simple task. In addition, quantities such as bending rigidity that are naturally defined on triangles would have to be properly mapped onto contacts lines between neighbouring cells. This is not straightforward to do. Developing a fully three-dimensional version of the AVM would be an even a greater challenge since the duality between Delaunay and Voronoi descriptions central to this model has no analogue in three dimensions.

We hope that the AVM will provide a useful and complementary tool for probing the aspects of the epithelial tissue mechanics that are not available to other methods, as well as serve as an independent validation for the results obtained by other methods.

## V. ACKNOWLEDGEMENTS

We would like to acknowledge many valuable discussions with Geoff Barton, Dapeng (Max) Bi, Luke Coburn, Amit Das, Tamal Das, Alexander Fletcher, MLisa Manning, MCristina Marchetti, Kirsten Martens, Inke Näthke, and K R Prathyusha. SH acknowledges hospitality of the Kavli Institute of Theoretical Physics during the workshop “From Genes to Growth and Form”. RS and CJW acknowledge support by the UK BBRSC (grant BB/N009789/1) and SH acknowledges support by the UK BBSRC (grant BB/N009150/1).

## APPENDIX I: FORCE ON THE CELL CENTRE

In this appendix we derive the expression for forces on the cell centre, Eq. (7), starting from the expression for the VM energy given in Eq. (1). The force on cell *i* is computed as the negative derivative with respect to **r***_i_* of the energy functional *E_VM_* in Eq. (1). Conceptually, this is a straightforward calculation with the only real complication being that *E_V M_* is most naturally written in terms of the positions of the Voronoi vertices, **r***_μ_*, while in the AVM we track positions of cell centres. In general,

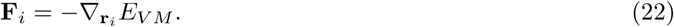

When computing the gradient in the last expression, we need to keep in mind that moving particle *i* changes the shapes of all of its neighbouring cells. Therefore, moving cell *i* exerts a force on a number of cell centres in its surrounding. All those contributions have to be taken into account when computing the force **F***_i_*. A direct consequence of this coupling between the cell and all of its neighbouring cells is that the force **F***_i_ cannot* be written as a simple sum of pairwise interactions between cell *i* and each of its neighbours. We’ll get back to this point below.

We start by computing the direct contributions resulting from moving particle *i* itself. The area term will produce a force

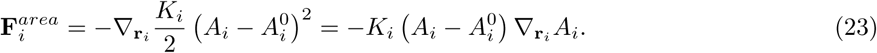

We need to compute 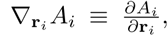 which we write by using the vector form of the chain rule for the calculating derivatives,

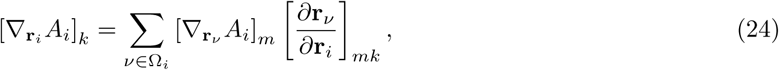

where the sum is over all vertices surrounding particle *i*, referred to as the *loop* Ω*_i_* of particle *i*. 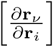 is the 3 × 3 Jacobian matrix of the transformation between coordinates of cell centres and positions of the vertices of the dual Voronoi tessellation. […]*_k_* represents the *k^th^*—component of the gradient vector, with *k* ∈ {*x, y, z*} and we assume summation over the repeated index *m*. The Jacobian can be computed using the barycentric coordinates that connect centres and vertices introduced in Eq. (5) and Fig. 2. We have

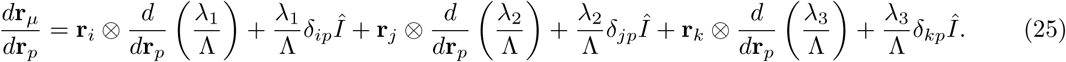

where

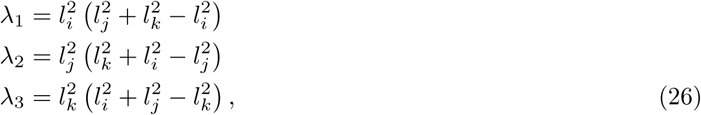

are the barycentric coordinates and Λ = λ_1_+ λ_2_+ λ_3_ with

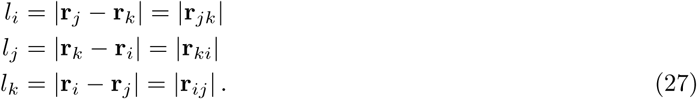

*Î* is a 3 × 3 identity matrix, and ⊗ stands for the outer product between two vectors. We proceed by calculating

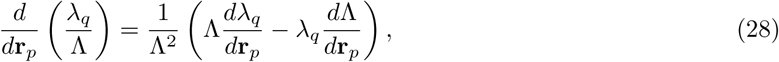

for *q* =1, 2, 3. It is straightforward to show that

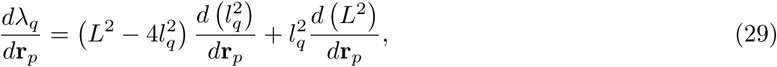

where *l_q_* is defined in Eq. (27) and 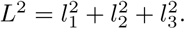 We readily calculate,

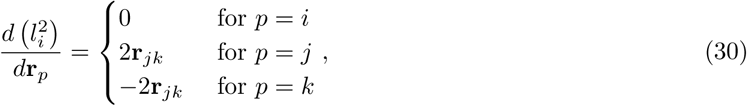

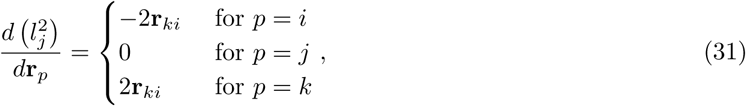

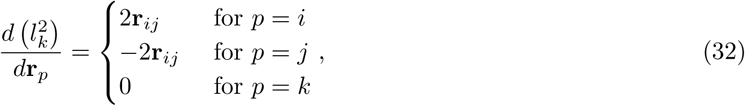

where **r***_ij_* = **r***_i_* – **r***_j_*, etc. Combining the last three expressions gives

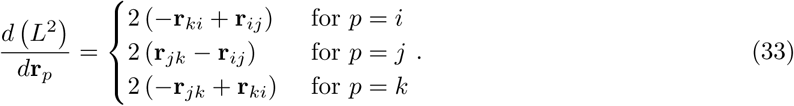

In order to reduce the computational effort, it is convenient to precompute and store derivatives for all three values of *p* = *i*, *j*, *k*. We start with the case *p* = *i* for which we have,

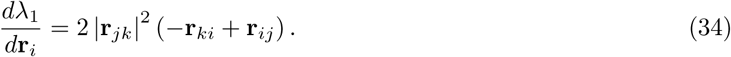

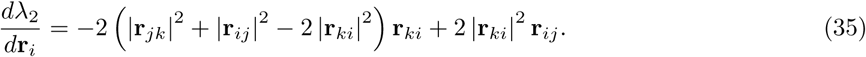

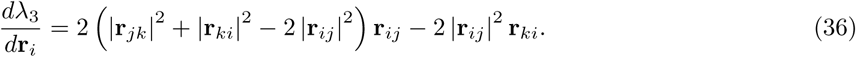

For *p* = *j* we have,

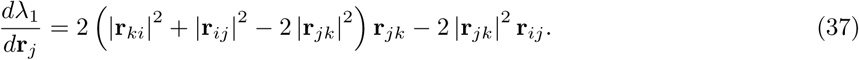

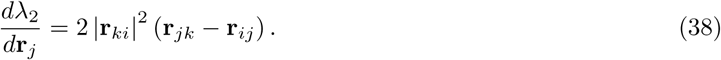

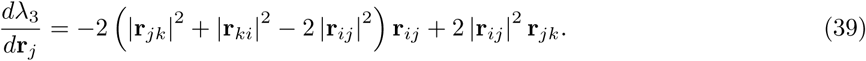

For *p* = *k* we have,

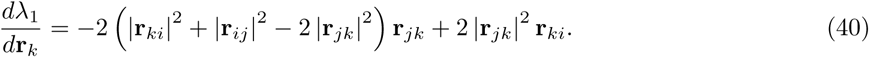

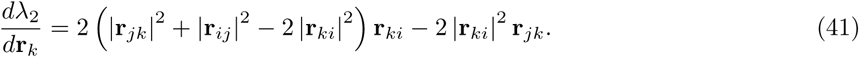

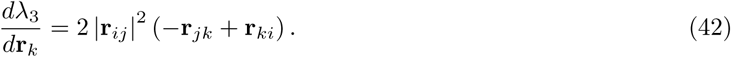

Finally,

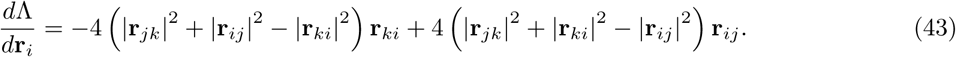

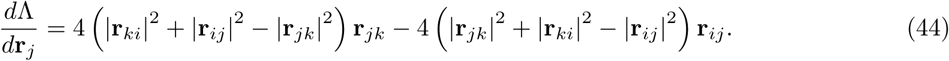

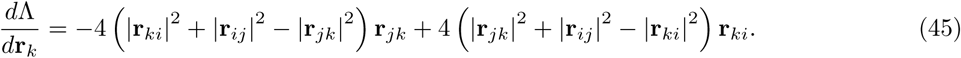

These expressions allow us to compute all derivatives in Eq. (28) and, in turn, the Jacobian in Eq. (25).

We still need to compute the derivative of the cell’s area with respect to the positions of the vertices of the Voronoi cell. A straightforward calculation starting from Eq. (3) gives

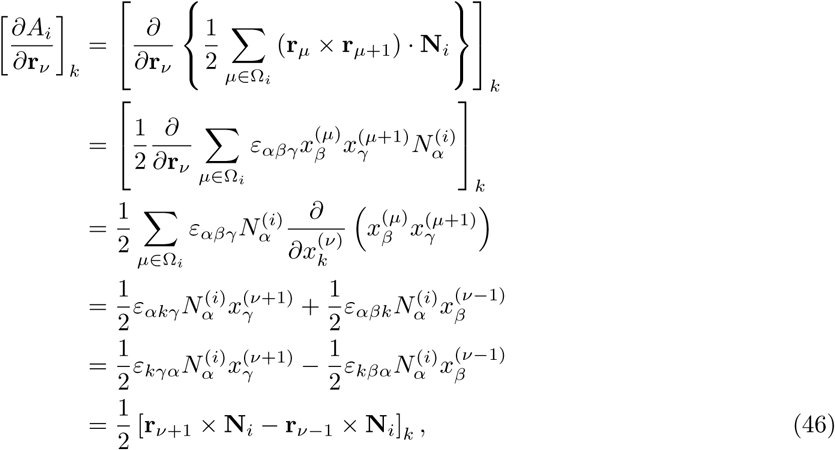
where *ε_αβγ_* is the Levi-Chivita symbol. Therefore, the expression for the area change due to displacing vertex *i* is

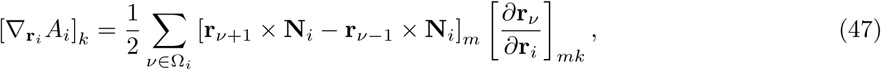
where, as above, we have assumed summation over the repeated index *m*. We finally derive the force on vertex *i* due to the area contractions,

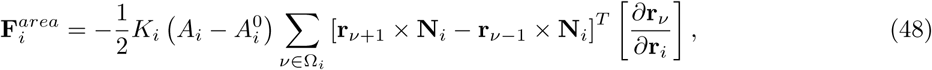
where *T* in superscript stands for transpose, i.e., […]^*T*^ […] stands for a matrix product between a 1 × 3 matrix (i.e., a vector) and 3 × 3 Jacobian matrix.

We can now proceed to calculate derivatives of the second term in Eq. (1). The perimeter of cell *i* is defined as

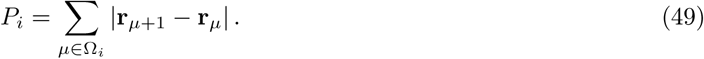

As above, we calculate

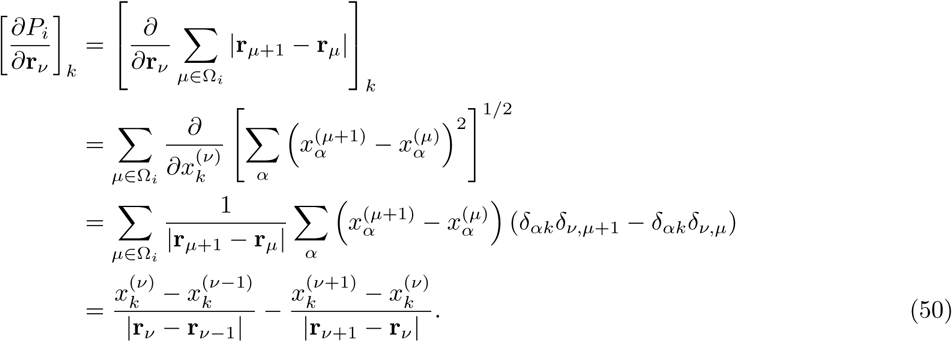

We therefore have

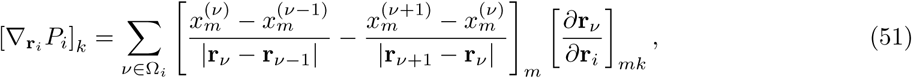
where we also assume summation over the repeated index *m*. The force term resulting from perimeter contractions is then given as

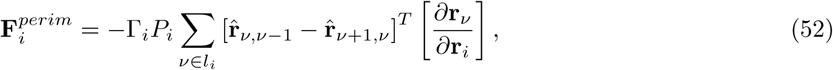

where we have defined 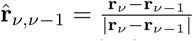, etc.

A similar calculation for the the last term in Eq. (1) leads to

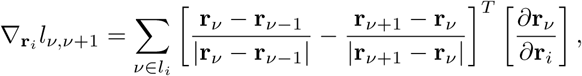

where we have explicitly labelled the two nearest neighbours (in the counterclockwise direction as) as *ν* and *ν* + 1. The force due to cell junction contractions is then computed as

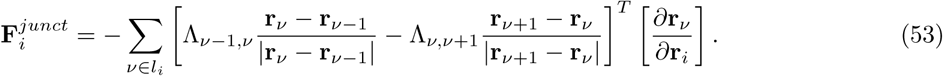

We now move to the second part, which is to determine the force on particle *i* as a result of displacing one of its surrounding particles. In order to do this, we go back to the original expression for the energy in the VM, Eq. (1). The force on vertex i is then

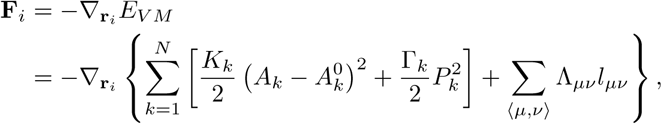

where *N* is the total number of cells. We first focus on the area and perimeter terms. We have

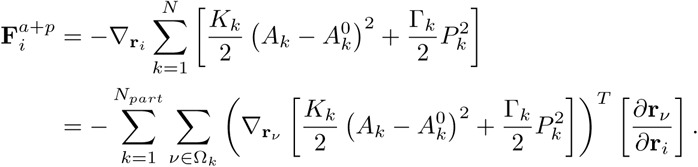

Further, we have

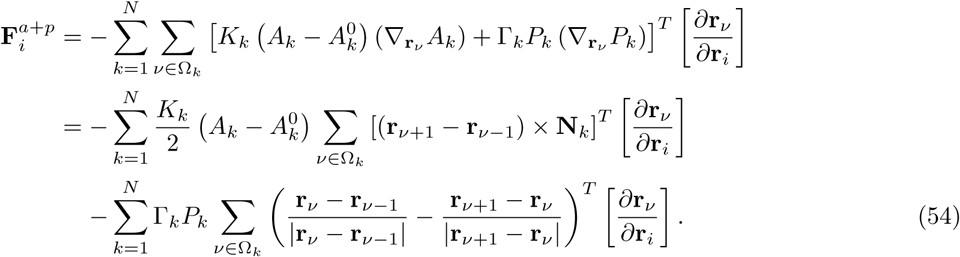

From the last expression it is clear that only vertices that are displaced by moving cell *i* are going to contribute to the force. These vertices are all “corners” of cell *i* and a subset of “corners” of its immediate neighbours affected by *i*. This gives us the algorithm for computing the total force on particle *i* coming from the area:

1. Loop over particle *i* and all its neighbours.

a. For particle *i* compute 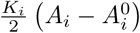 and multiply it by the sum 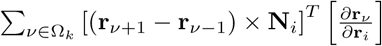. Note that this sum is over all vertices (corners) *ν* of the cell *i*.
b. For all immediate neighbours *j* of cell *i* compute 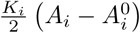 and multiply it with the sum 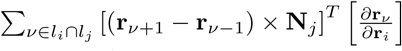. Note that *ν* ∈ *l_i_* ∩ *l_j_* ensures that vertices *ν* surrounding *j* are taken into account only if they are affected by (and also belong to) cell *i*.

A similar algorithm can be used to compute force contribution of the perimeter term.

We now focus on the last term, which is the force along the cell junctions. If we note that we can write the cell junction term as

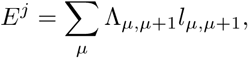

we have

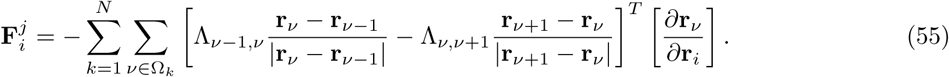

As before, we loop over all vertices that are affected by changing the position of cell *i*, which are all corners of the cell *i* and a subset of corners of its immediate neighbours whose positions are determined by **r***_i_*.

Combining Eqs. (54) and (55) then ultimately leads to Eq. (7). We note that in Eq. (55) we have absorbed factor of 1/2 arising from double counting edges into the definition of Λ_*ν,ν*+1_.

Finally, we briefly address another contribution to the force which needs to be added to the model allow for simulations deep in the liquid-like phase. As discussed in Sec. IIIB, similar to the SPV model, the AVM shows a transition between solid-like and fluid-like phases. In the solid-like phase, intercalation events are inhibited and without cell division and death, cells do not exchange their neighbours. In the fluid-like phase, on the other hand, cells are much more mobile and one observes a large number of T1 transitions and intercalations. In this regime, the Delaunay triangulation is very irregular with many obtuse triangles. If a triangle becomes very obtuse, its circumcenter is far outside the triangle and even small changes in the position of one of its vertices leads to large movements of the circumcenter. For any simulation time step that is not extremely small, this can lead to unphysical self intersections of the triangulation and cause the simulation to be unreliable, or, in the most extreme cases, causes it to crash. In practice, in order to prevent this from happening while still being able to use a reasonably large time step, we endow each cell centre with a soft repulsive core of radius *a*. The repulsive potential between neighbouring cell centres is then given as

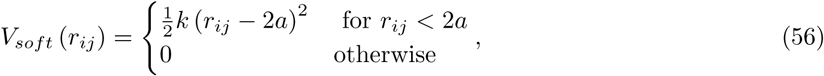

where *r_ij_* = |**r***_i_* — **r***_j_*|. This core prevents two cell centres from getting too close to each other. The corresponding force on cell *i* is

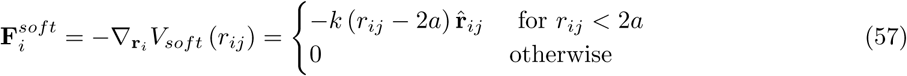

with r̂*_ij_* ≡ |**r***_i_* — **r***_j_*| /*r_ij_* and according to the Newton’s third law, 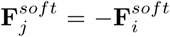. The soft repulsion does not interfere with the AVM dynamics except for its regularising effect. In Fig. 6e-g, the pale spheres drawn at the cell centres have radius *a*, and they remain far removed from the cell boundaries.

We note that this repulsive force is similar in spirit to the limits that have to be imposed on edge lengths in order to prevent unphysical self-intersections in the triangulated models for lipid membranes, that have been extensively studied in the 1990s.^99^ It is interesting to note that in the case of triangulated models for lipid membranes one also needs to introduce a maximum allowed edge length in order to prevent unphysical configurations. In the case of AVM, this is not necessary.

## APPENDIX II: ALGORITHM FOR HANDLING BOUNDARIES

Due to the dynamic nature of the model, even without cell division and death, it is not possible to retain a constant number of boundary particles. Instead, the boundary line has to be able to contract or extend in order to accommodate changes inside the tissue. This is achieved by dynamically adding and removing boundary particles.

We first focus on the boundary expansion. We require that all cells are contained within the boundary, that is, no dual vertices belonging to a cell are allowed to “spill” over the boundary line. This condition is violated if the angle opposite to a boundary edge is greater than 90°. In this case, the centre of the circumscribed circle falls outside the triangle and, therefore, outside the boundary. In order to prevent this from happening we perform the following check (see also Fig. 12):

1. For each boundary edge *e* compute angle *α_e_* at the particle *p_e_* opposite to it.
2. If *α* > 90^°^
  a. Compute the position, **r***_Pn_*, of the new particle *p_n_* by mirroring the coordinates of *p_e_*, **r***_pe_*, with respect to edge *e*. If **r̂***_e_* is the unit-length vector along edge *e* then

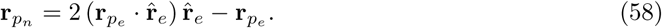
  b. Add a new boundary particle *p_n_* at position **r***_pn_* and mark it as boundary.
  c. Remove boundary edge *e*, i.e., the two boundary particles at its end are no longer neighbours.
  d. Connect *p_n_* to the two boundary particles disconnected in (c).
  e. Connect *p_n_* to *p_e_*.
3. If at least one new boundary particle was added in 2., rebuild the triangulation.

**Figure 12.**
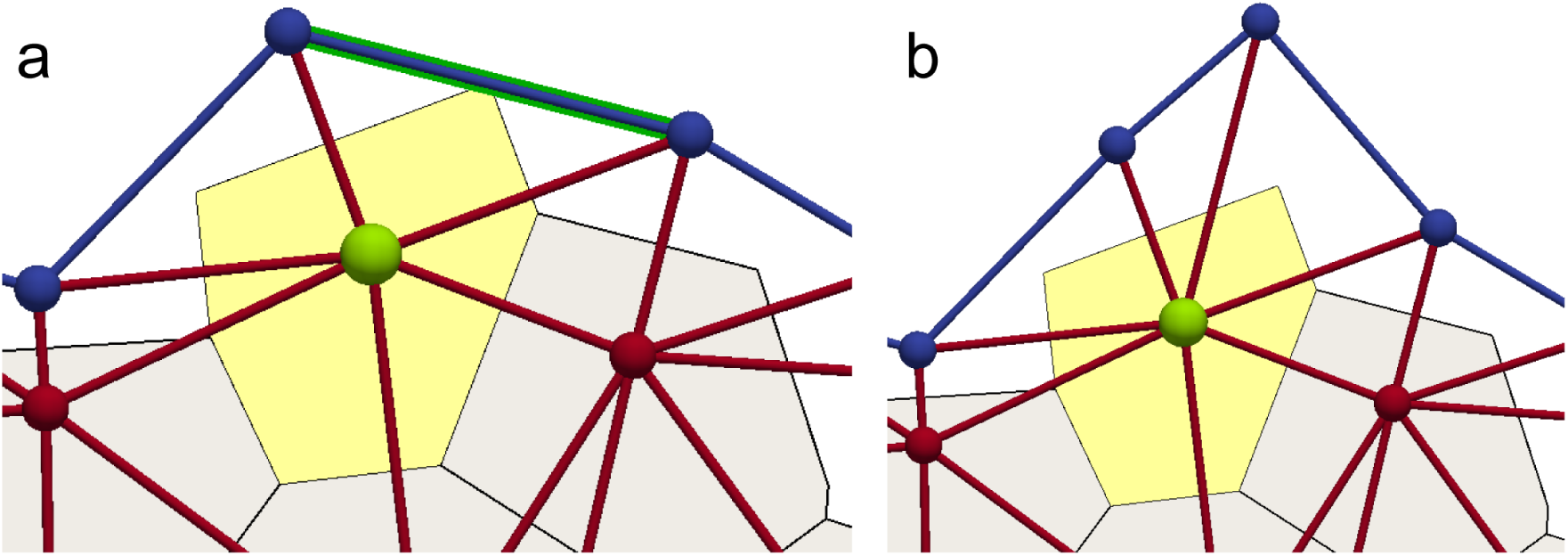
Expansion of the boundary by adding a new boundary particle. (a) If the angle at the internal particle shaded in green opposite to the highlighted edge reaches 90°, one of the corners of the cell (yellow polygon) touches the highlighted edge. This triggers a “flip” mechanism. (b) The internal particle shaded in green is mirrored along the shaded edge in (a) and a new boundary particle (top blue) is introduced. The shaded edge is flipped such that the new particle is connected to the “green” one.

Note that the procedure outlined above always converges in a single step.

Shrinking of the boundary is achieved by removing particles that have no connections to the internal particles. In this situation, no part of a cell can be inside a triangle that has two of its sides being boundary edges and it can be safely removed. The algorithm schematically outlined in Fig. 13 is as follows:

1. For each boundary particle p compute number of edges *n_e_* (*p*) that the particle belongs to.
2. If *n_e_* (*p*) ≤ 2 remove *p*.

**Figure 13.**
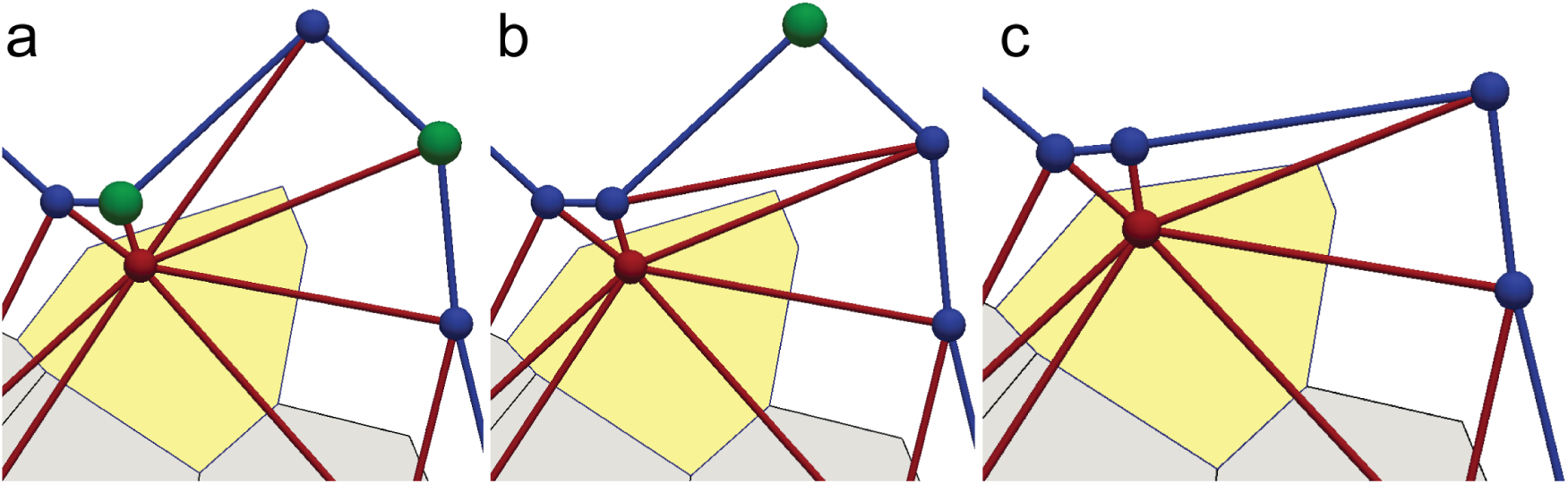
Shrinking of the boundary is achieved by removing boundary vertices that have only two bonds. (a) If the sum of angles at two particles shaded in green opposite the edge connecting the internal and boundary particles is greater than 180° the edge is flipped (this is the standard equiangulation move that occurs for all internal edges). (b) After the flip, the boundary particles shaded in green has only two bonds, both to its boundary neighbours. (c) The “green” particle in (b) is then removed.

Note that the position of the “dangling” particles does not directly affect the shape of the cell.

An important point to make here is that the algorithms used to add and remove boundary particles *do not* lead to sudden discontinuous changes of the shape of the cell or the force acting on its centre. However, addition and removal of boundary particles inevitably leads to discontinuous changes in the forces acting on boundary particles. A potential way to avoid such discontinuous behaviour would be to, e.g., smoothly “turn on” the interactions with newly added particles or by slowly “fade out” interaction with particles that are to be removed. In practice, however, the discontinuous causes by simply adding or removing boundary particles lead to changes in the force that are small and do not appreciably affect the simulation.

## APPENDIX III: IMPLEMENTATION

The AVM is implemented into the *SAMoS* code developed by this team.^100^In this appendix we first provide a general overview of the organisation of the *SAMoS* code and then discuss how the AVM in implemented in it.

### SAMoS overview

*SAMoS* is a software package developed for simulating agent-based active matter systems confined to move on curved or flat surfaces.^74^ The code is written in C++, using the C++98 standard with extensive use of the Standard Template Library and boost libraries.^101^ It utilises a modular, object oriented design making it very flexible and simple to extend. It adheres to modern software design principles and uses a cross-platform build system *(cmake*) as well as automatic documentation generation with *Doxygen.^102^*

*SAMoS* consists of several components each implemented as class hierarchies.

- **System** - central component that handles the system configuration;
- **Parser** - a recursive descent parser for parsing files that control execution of the simulation (configuration files). It is implemented using boost’s Phoenix library;^103^
- **Messenger** - logs system messages (warnings, errors, etc.) as well as meta-data (parameter trees in JSON or XML format) for data curation;
- **Neighbour list** - handles build and update of the Verlet neighbour list;^64^
- **Constraint** - handles projection on various flat or curved surfaces. It is possible to have multiple constraints acting on different groups of particles;
- **Interactions** - handles all interactions on and between particles. Interaction parameters can be type-specific, i.e., it is possible to simulate multicomponent systems. It is also possible to have multiple interaction types simultaneously present in the system;
  - **Pair/multi-body** interactions - handles all interactions that involve pairs or multiplets of particles (Vertex, Lennard-Jones, soft repulsion, Morse, etc.);
  - **External** interactions - handles all forces that act on a single particle, such as external fields and/or activity;
  - **Bond/angle** interactions - handles interaction between connected beads for simulating filaments;
- **Alignment** - handles alignment of the particle orientation
  - **Pair** alignment - handles various models for alignment to the direction of neighbouring particles (discussed in Sec. IIB 2);
  - **External** alignment - handles alignment to internal or external cues, such as velocity, cell shape or external fields (discussed in Sec. IIB2);
- **Dump** - handles output of the system snapshot in various formats (VTP, raw text data file, input configuration for restarts, etc.). It is possible to have multiple dump types present in the same simulation;
- **Log** - handles output of the current system state, such as total potential energy, mean velocity, temperature, etc.;
- **Population** - handles addition and removal of particles, such as during cell division and death. It is possible to have multiple population controls acting at the same time or on different groups of particles;
- **Integrator** - handles various numerical integrators (Langevin, Brownian, NVE, etc.) for solving the equations of motion. Different integrators can acts on different groups of particles (e.g. no motion, for keeping a subset of particles fixed).

The components are designed to be as loosely coupled as possible, in order to ensure flexibility, ease of testing and debugging; and also to make extensions of the code, such as adding a new interaction force or population control mechanism, as simple as implementing a new subclass with a minimal need to modify the core of the code.

In order to perform a simulation with *SAMoS,* the user has to supply two or three text files: 1) a file, referred to as the *data* file, containing the initial configuration of the system (i.e the initial positions and velocities of the particles, particle types, polarity, etc.), 2) for AVM simulations only, a file, referred to as the *boundary* file, containing the initial labels and connectivity of the boundary particles and 3) the parameter file, referred to as the *configuration* file, which sets the simulation protocol (interaction types and parameters, constraints, type and frequency of dumps, simulation time step, etc.). Commands in the configuration file are parsed and executed in the order they appear. Examples of data and configuration files can be found in the *configurations* directory in the *SAMoS* installation.

In the current implementation, *SAMoS* runs on a single CPU core.

### AVM implementation

The AVM is implemented as an extension of the *SAMoS* code. The main addition to the code involves a light-weight implementation of the half-edge data structure^104^ as a separate, *Mesh* class. This class holds the information about the Delaunay triangulation and computes its dual Voronoi diagram. The *Mesh* class also ensures that the Delaunay character of the triangulation is preserved between rebuilds using the equiangulation procedure discussed in Sec. IIB 5. Finally, *Mesh* computes the Jacobian matrix,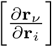and supplies its elements to the part of the code that computes the force on the cells. A feedback loop from the integrator ensures that the *Mesh* class always has the correct position of the cell centres.

The Delaunay triangulation of the initial positions of the cell centres is performed using CGAL’s Delaunay library.^105^ In order to properly include the boundary of the cell sheet, we compute a constrained Delaunay triangulation,^106^where the boundary line, supplied by the user as an input file, acts as the constraint placed on the triangulation. It is important to note that all Delaunay triangulation libraries always produce a convex hull of the region that needs to be triangulated. The existence of the boundary line and the use of the constrained triangulation method allows us to simply and clearly distinguish between the triangles that are “inside” the tissue and should be kept and the “outside” ones that are an artefact of the triangulation procedure and should be discarded. This weeding out of outside triangles imposes some performance burden on the computation and we do not perform it at every time step, but maintain it to be Delaunay by applying equiangulation moves that are fast to compute. The triangulation is completely rebuilt only after the steps that involve cell division/death events or boundary extension/contraction. In order not to disturb the connectivity of the boundary, only the internal edges can be flipped. In practice, depending on the parameter values, the triangulation is rebuilt once every 10-50 time steps. This substantially improves the performance of the code.

The force on the cell centre, Eq. (7), is computed by a subclass of the pair/multi-body submodule in the Interactions module discussed in the previous section. This class invokes the *Mesh* class in order to obtain the information about positions of dual vertices and components of the Jacobian matrix.

An overview of the general organisation of the AVM implementation is shown in Fig. 14.

**Figure 14.**
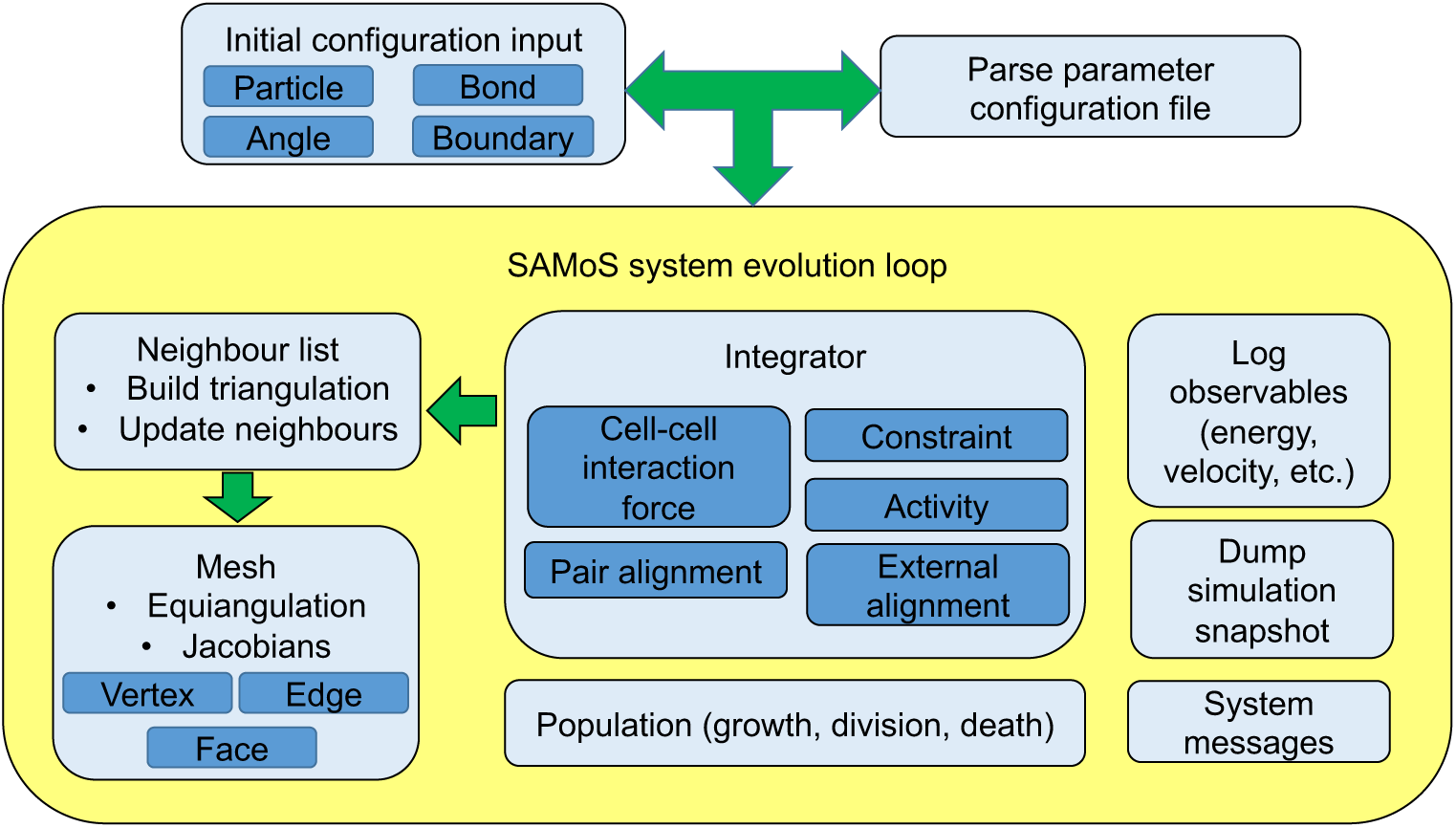
Overview of the general organisation of the AVM implementation into *SAMoS*.

